# A conserved phenylalanine motif among Teleost fish provides insight for improving electromagnetic perception

**DOI:** 10.1101/2024.04.04.588096

**Authors:** Brianna Ricker, E. Alejandro Castellanos Franco, Gustavo de los Campos, Galit Pelled, Assaf A. Gilad

## Abstract

Magnetoreceptive biology as a field remains relatively obscure; compared to the breadth of species believed to sense magnetic fields, it remains under-studied. Here, we present grounds for the expansion of magnetoreception studies among Teleosts. We begin with the electromagnetic perceptive gene (EPG) from *Kryptopterus vitreolus* and expand to identify 72 Teleosts with homologous proteins containing a conserved three-phenylalanine (3F) motif. Phylogenetic analysis provides insight as to how EPG may have evolved over time, and indicates that certain clades may have experienced a loss of function driven by different fitness pressures. One potential factor is water type with freshwater fish significantly more likely to possess the functional motif version (FFF), and saltwater fish to have the non-functional variant (FXF). It was also revealed that when the 3F motif from the homolog of *Brachyhypopomus gauderio* (B.g.) is inserted into EPG – EPG(B.g.) – the response (as indicated by increased intracellular calcium) is faster. This indicates that EPG has the potential to be engineered to improve upon its response and increase its utility to be used as a controller for specific outcomes.

## Introduction

The ability to sense and perceive the magnetic field of the Earth is becoming a widely discussed idea pertaining to an abundance of species (*1-4*). The best-known examples include migratory species that travel considerable distances throughout their lifetimes for optimal living and breeding conditions. One such example is the European robin (*Erithacus rubecula)* – a migratory bird that has been at the forefront of magnetoreception research after the proposal of the radical pair mechanism relying on retinal Cryptochrome 4 (Cry4) (*5*).

Salmonids of the genus *Oncorhynchus* have also been common subjects of magnetoreception studies due to their intrinsic natal homing (*6*). Pink salmon (*Oncorhynchus gorbuscha)* have demonstrated magnetoreceptive abilities that are essential for them to migrate around the Pacific and back to their native rivers to spawn (*7*). Although some studies indicate that salmonids may use magnetite – iron containing crystals – to sense the Earth’s magnetic field (*8*), this theory remains open for debate (*9-11*).

Many non-migratory species have also been shown to have magnetoreceptive properties; one of particular interest being the glass catfish (*Kryptopterus vitreolus*) (*12*). This is a small catfish (up to 6.5cm in length) that resides in slow moving, murky streams in Thailand (*13*). Previous studies have indicated that the glass catfish retreats away from electromagnetic stimulus, and is believed to be facilitated by the electromagnetic perceptive gene (EPG) protein (*12, 14*).

EPG is a small (∼9kD) protein that adopts a Ly6/UPAR three-finger structure and is membrane associated via glycosylphosphatidylinositol (GPI) anchor (*15*). EPG has shown response to magnetic stimuli in mammalian cells in the form of increased intracellular calcium (*12, 15-17*), in activatable split protein systems (*18*), and in rat models with increased neuronal plasticity (*19*) and reduced seizure activity (*20*), but its mechanism of action remains elusive. Site-directed mutagenesis has indicated a region of interest – the three-phenylalanine (3F) motif – in the protein that is essential for its function (*15*), but it remains unclear how this region facilitates magnetoreception.

There exists value in EPG for the field of synthetic biology as a potential method of external control of molecular devices and synthetic circuits. However, its current state of being poorly understood makes it difficult to use and implement such technologies. Therefore, this study aims to expand upon the bank of knowledge surrounding EPG to make future expansion of magnetoreceptive biology and synthetic technologies using this property more probable. Understanding the origins of the protein may not only provide insight into its intrinsic use in fish but also serve to guide future efforts to improve upon the function of EPG and make it more applicable for specific uses.

## Results & Discussion

### Translated Basic Local Alignment of EPG

The NCBI’s tblastn tool was used to identify homologous proteins to EPG. This tool searches a specified database for nucleotide translations of the input amino acid sequence (*21*). In this case, tblastn returned 62 unique species that express a protein with high sequence homology to EPG. An additional 10 species with EPG homologs in their genomes were either discovered in the EFISH Genomics database, or uncovered by manually searching unannotated genomes within NCBI databases of species in the same genus as others with an EPG homolog.

In total, 72 species were discovered to have a homolog of EPG. In Figure 1A, the sequences of these homologs are displayed in alignment with the sequence for EPG. Visually, high homology between the sequences can be observed. This alignment also highlights the conservation of the previously discovered 3F motif – highlighted in blue. The phenylalanine (F) residues in this region are known to be critical in order for EPG to sense and respond to magnetic fields (*15*). Figure 1B depicts a sequence logo of this highlighted section of the alignment. It becomes clear that the F in positions 1 and 10 are highly conserved, but the F in position 7 has some variability. As discussed in a previous article, mutating this F residue results in a loss of function in EPG (*15*). Therefore, species that adopt the FXF motif may be less likely to have magnetoreceptive abilities associated with EPG, while FFF species may be more likely.

**Figure 1.**
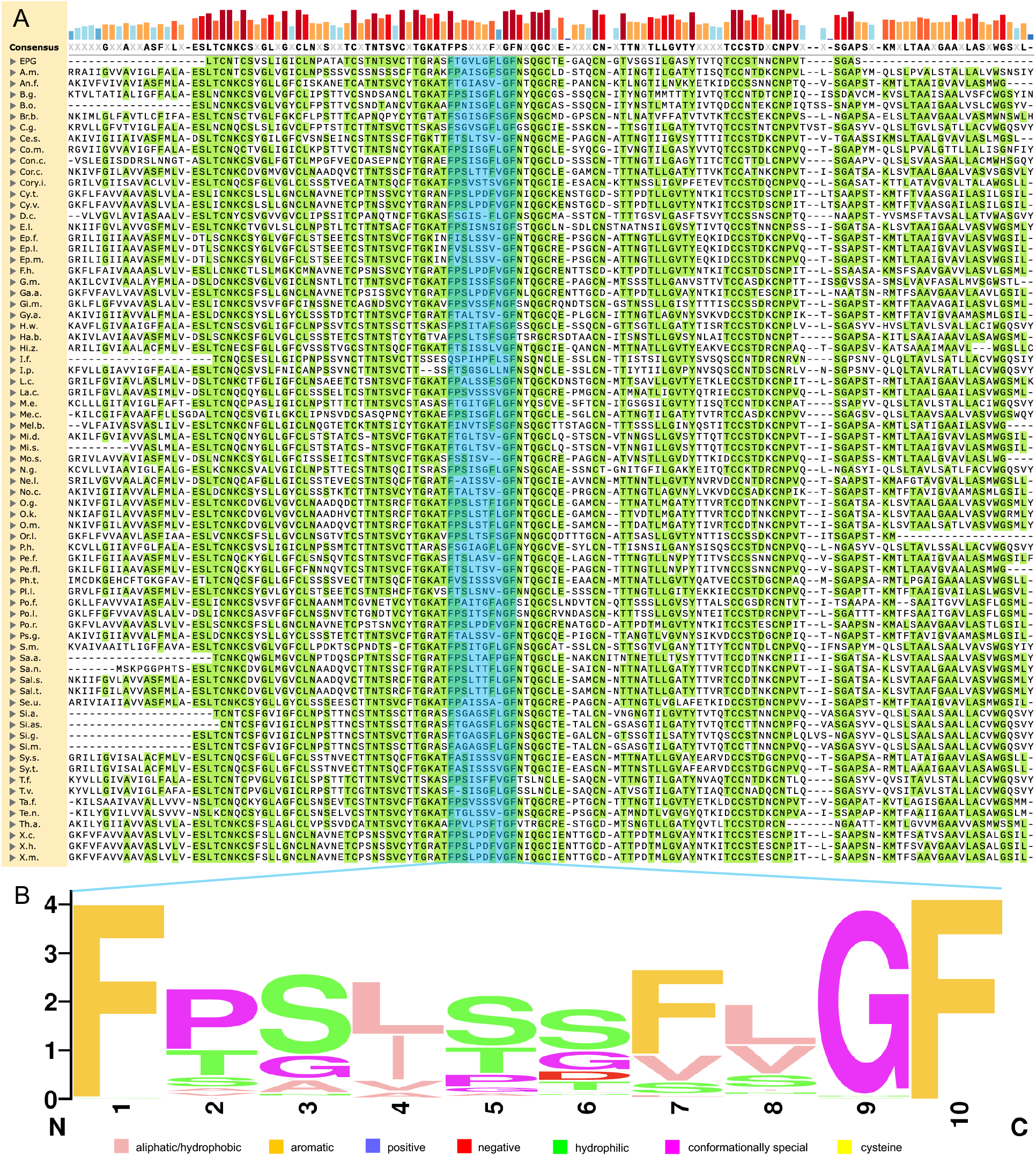
Homologs of EPG are revealed with tblastn. (A) MUSCLE amino acid alignment of EPG with homologs discovered with NCBI’s tblastn tool. Amino acids highlighted in green match the consensus sequence. The 3F motif is highlighted in blue. The height and color depth above each amino acid indicate how conserved that particular residue is across all the sequences. The alignment was created using SnapGene. (B) The sequence logo of the 3F motif exhibits the most common amino acid, as well as the frequency of amino acids in each position in the motif. Amino acids are labeled with the Zappo color scheme to reflect their physical chemical properties. The sequence logo was generated using WebLogo (*22*).

### Phylogenetic Assessment of Species with an EPG Homolog

To assess how these 72 fish species are related to the *Kryptopterus vitreolus* and each other, a phylogenetic tree was constructed using TimeTree (*23*), as shown in Figure 2. We observe that all species that exhibit EPG or a homolog belong to the Teleostei infraclass. Clear divisions between FFF and FXF species are also made apparent; paired with the idea that FFF fish may be magnetoreceptive and FXF fish may not, we ask whether this trait is gained or lost over the course of this phylogeny.

**Figure 2.**
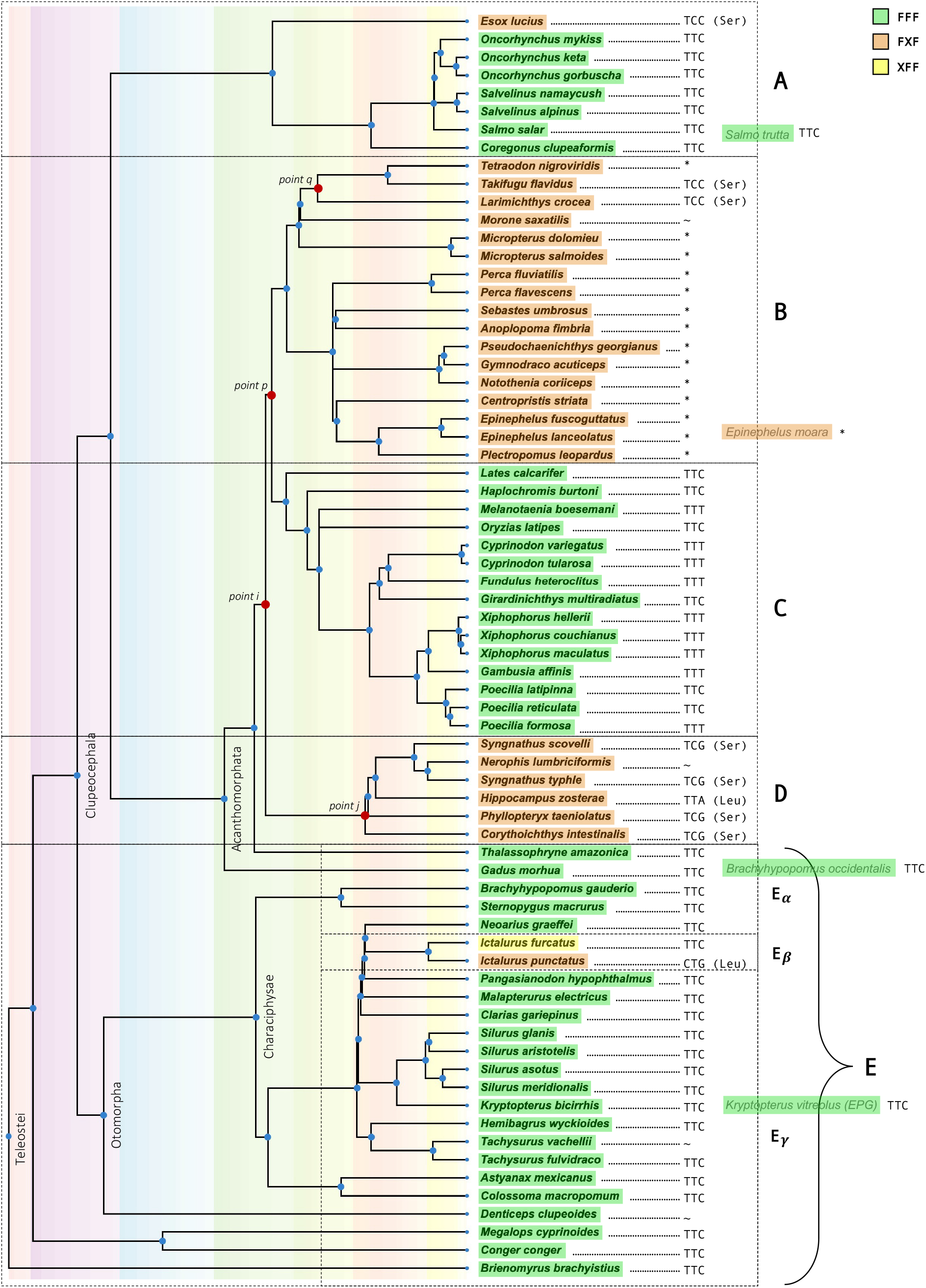
Phylogenetic tree of all the species that have an EPG homolog discovered by tblastn. The tree was generated using TimeTree (*23*). Four species were unresolved by TimeTree (right shifted in gray text) but their approximate positions within the tree have been inferred by their genus. Species highlighted in green contain all three F residues within their 3F motif (positions 1, 7, and 10). Species highlighted in orange do not have F in position 7. Highlighted in yellow is one species that differs without the F in position 1. To the right of each species is the codon used for the amino acid in position 7. Codons, other than those for F, also show their respective translation. The star (^*^) represents sequences that have had a 3bp deletion of nucleotides 15-17. The Tilda (∼) represents sequences that have had other deletions of 3bp or 6bp. The tree has been separated into several pieces to facilitate discussion.

Analysis of the DNA sequences for these 3F motifs provides a more defined picture that may help us answer the question at hand. In group B, based on *Takifugu falvidus* (Ta.f.), and *Larimichthys crocea* (La.c.), we can infer that there was a simple point mutation from TTC → TCC [F → S] at point *p*. This mutation may have led to a loss of function of the protein, which effectively removes selective pressure and further opens the possibility of more mutations to happen at a faster rate – such as the deletions observed throughout the rest of group B. Point *q* then represents the continued lineage of fish that have the original point mutation versus the deletion adopted by the rest of group B.

The mutations that form group D could have occurred at either point *i* or point *j* with the latter more probable. A mutation at point *i* from TTC → TCC [F → S] would then need to undergo a reverse mutation to be consistent with group C, *i*.*e*. TTC → TCC → TTC [F → S → F]. Although this is possible, a much more likely scenario is that there was a point mutation that occurred at point *j* that led to the formation of group D (Table 1).

**Table 1:** Evolutionary changes in the codon that encodes position seven of the 3F motif.

|  |  |  |  |
| --- | --- | --- | --- |
| **No mutation** | TTC (Phe) |  |  |
| **1<sup>st</sup> mutation** | TCC (Ser) | TTG (Leu) | TTA (Leu) |
| **2<sup>nd</sup> mutation** | TCG (Ser) |  |  |

Group E_β_ also points to a loss of function over the phylogeny. *Ictalurus punctatus* (I.p.) contains two mutations (TTC → TT<u>G</u> → <u>C</u>T<u>G</u> [F → L]) likely rendering the protein nonfunctional as it harbors the FXF motif. *Ictalurus furcatus* (I.f.) differs from every other species though and is missing the first F in the motif instead of the F in position 7. A previous study indicates that mutating the first F in EPG’s 3F motif (to obtain a XFF variant) did not lead to a loss of function (*15*), and therefore may be true of the I.f. homolog.

With this information at play, the most likely scenario is that early Teleost fish exhibited EPG, but the function was lost as the group of fish diversified. This conclusion also remains consistent with the general trends of the Teleost fish. A significant increase in anatomical and physiological differences is observed among Teleostei after the Cretaceous–Paleogene (K–P) extinction (*24*). Today, the Teleostei are the largest group of fish on the planet, containing approximately 95% of all living fish species, exhibiting incredible biodiversity and inhabiting waters across the globe (*25*). The Teleostei are currently being studied for numerous unique attributes including electrical abilities (*26-28*), mouthbrooding (*29*), brackish tolerance (*30*), cold tolerance (*31, 32*), among others, and now including magnetoreception.

### Phenotyping Teleostei to Analyze Motif Variants

Because these 72 fish species differ so much in their ecological adaptation and in morphology, we sought to link their characteristics to their motif type to uncover if fish with the FFF motif follow any trend that could tell us more about the native purpose of the protein and what these fish might use magnetoreception to accomplish.

The most obvious place to start is with migratory habits since animals are widely believed to sense the Earth’s magnetic field to navigate. In Figure 3A we observe both motif types have migratory and non-migratory species. The association between the presence of the FFF motif and migratory habits was statistically significant (p-value 0.005, Figure S1 and Table S1); however, this result is driven by a large number of species with undetermined migratory behavior and a significantly lower frequency of the FFF motif among those species compared to migratory and non-migratory ones (Table S1); thus, making the data indiscernible at this point.

**Figure 3.**
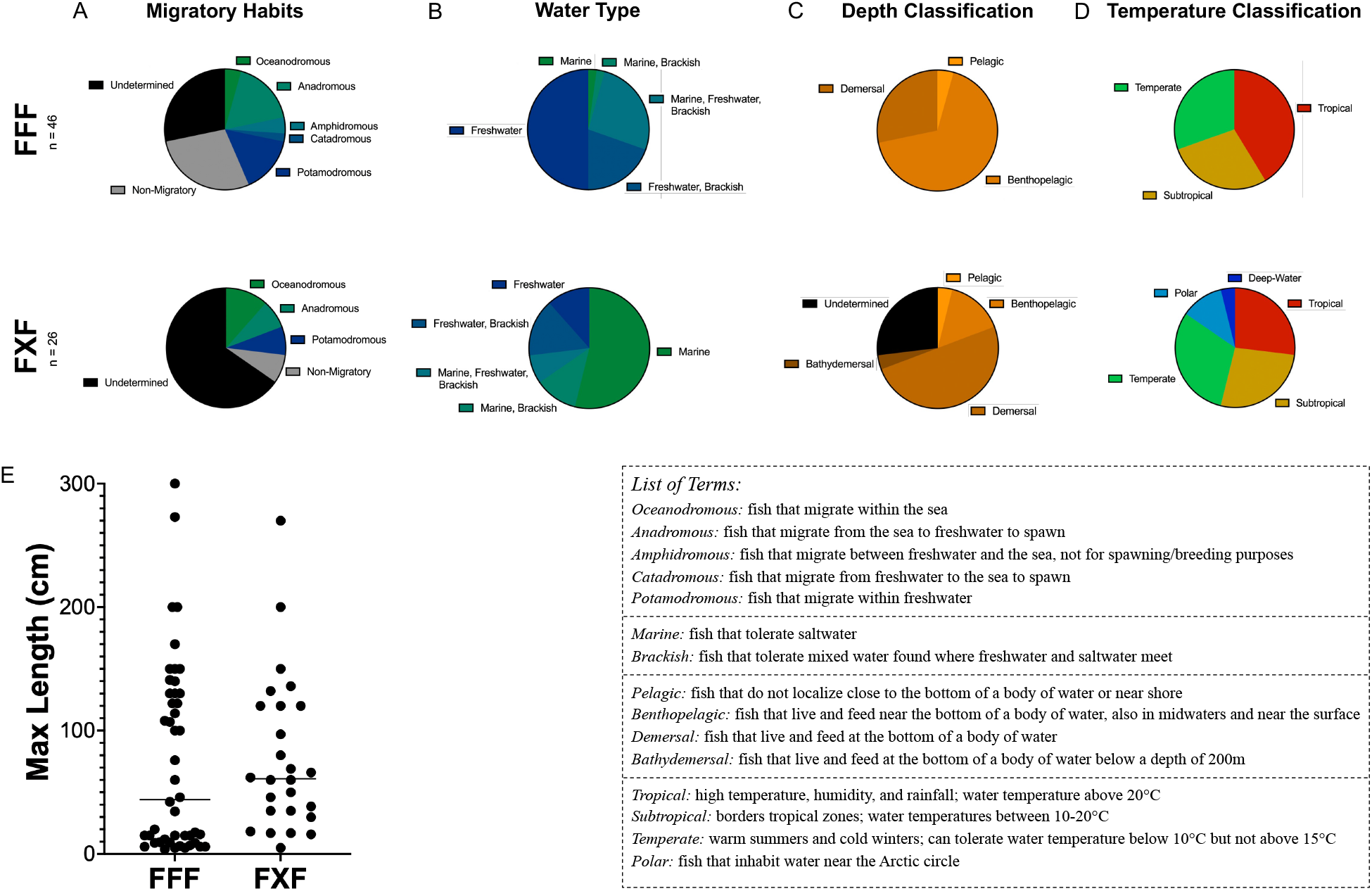
Characteristics of fish species with an EPG homolog separated by FFF and FXF 3F motif types. (A) Migratory habits of FFF vs. FXF species. (B) Water tolerance of FFF vs. FXF species. (C) Depth classification of FFF vs. FXF species. (D) Temperature classification of FFF vs. FXF species. (E) Maximum recorded length of each species grouped by FFF and FXF. Characteristics for each fish were gathered from FishBase (*33*). (See Figures S1 through S4 and Tables S1 through S4 for results from statistical tests for the association between the presence of the FFF motif and species’ characteristics).

Figures 3B details the type of water these fish inhabit; we found a statistically significant association between water type and the presence of the FFF motif (Figure S2). Interestingly, fish with the FFF motif are significantly more inclined to freshwater, whereas FXF fish are more inclined to saltwater. Of the solely freshwater fish, ∼88% (+/−6.2%) have the FFF motif and of the solely marine fish, ∼93% (+/−6.4%) have the FXF motif. Although the association between water type and the presence of the FFF motif was highly significant (Figure S2 and Table S2), we do not know why the FFF variant would be specifically advantageous to fish in freshwater environments.

Another characteristic we examined is the depth classification of these fish as shown in Figures 3C and S3. Most of the species we studied, both FFF and FXF, are benthopelagic or demersal. In general, these fish tend to be found near the bottom of a body of water, and sometimes these terms are used interchangeably with the slight difference being that benthopelagic fish are sometimes also observed in mid-to surface waters. We found a highly significant (p-value<0.001, Table S3) association between water depth and the presence of the FFF motif, with benthopelagic species having a much higher frequency (88.6% +/−5.4%) of the FFF motif relative to demersal species (50% +/−9.8%, Figure S3). However, the introspective nature of these terms leaves some subjectivity in the significance observed.

In Figures 3D and S4, we show the temperature classification of the fish. Although species that live in tropical water had a slightly higher prevalence of the FFF motif (73.1%, +/−8.7%, Figure S4) compared with those living in subtropical (65.0% +/−10.7%) or temperate environments (63.6% +/−10.3%), the association between the presence of the FFF motif and water temperature was not statistically significant once we accounted for multiple testing (p-value 0.027, Figure S4). These findings reiterate that the Teleostei inhabit nearly every corner of the globe.

Lastly, in Figure 3E, we show the maximum recorded length of each species as a metric for the size of the fish. The thought process here was that the FFF motif may have been advantageous for smaller prey fish as a means of detecting a shift in the magnetic field as a predator approaches, ultimately causing them to flee. On average, species carrying the FFF motif were 2.3cm shorter than those with FXF motif (Tables S4 & S5); however, this difference was not statistically significant (p-value 0.895). It is worth noting that the distribution of body length among species carrying the FFF motif appears to be bimodal, with an apparent enrichment for short species and another group having longer body size (Figures 3E & S6).

### Mutagenesis of EPG’s 3F Motif Informed by Homologs

Another important prospect towards understanding the diversity within EPG was to examine how different 3F motifs function. Because we previously identified that the 3F motif is critical for EPG to function, we sought to mutate the 3F motifs from these other species into EPG from the *Kryptopterus vitreolus*. The 3F motif of EPG was replaced with the 3F motif of homologs of a few closely related fish from the Characiphysae superorder (*Brachyhypopomus gauderio* (B.g.), *Ictalurus punctatus* (I.p.), *Sternopygus macrurus* (S.m.), and *Malapterurus electricus* (M.e.)) which are displayed in Figure 4A. These EPG mutants were then subjected to a previously described assay in HeLa cells with GCaMP6m (*15*) to observe changes in intracellular calcium. Figure 4B shows the native function of EPG in mammalian cells – intracellular calcium increases upon magnetic stimulation.

**Figure 4.**
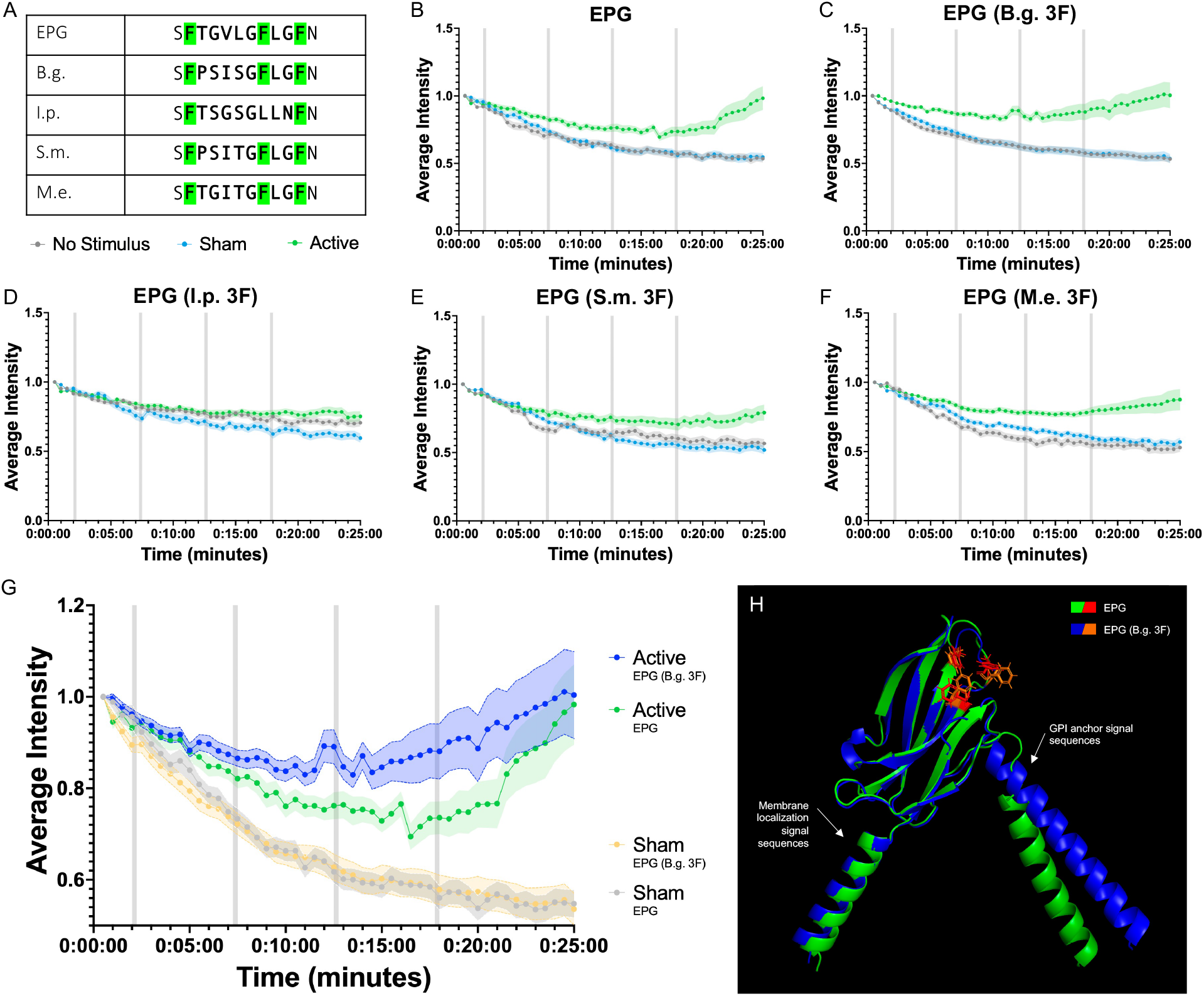
Utilizing a GCaMP6m assay to compare the efficacy of 3F motifs from homologs when inserted into EPG. (A) Comparison of 3F motifs from EPG and several homologs from closely related members of the Characiphysae superorder. (B-F) HeLa cells expressing versions of EPG-IRES-GCaMP6m subjected to a GCaMP6m functional assay; cells were exposed to either active, sham, or no stimulus with a pulse pattern of 15s on 5min off for 4 pulses (gray bars). Error bars are representative of 95% CI. Experiments include n=90 cells over three experiments for no stimulus, sham, and active groups respectively. (B) EPG-IRES-GCaMP6m. (C-F) EPG 3F homolog mutants. (G) Data from EPG and EPG (B.g. 3F) displayed together to highlight their significance. (H) Predicted structure of EPG (B.g. 3F) aligned with the predicted structure of EPG created using PyMOL (*34*).

Figures 4C-F show the data for the EPG mutants. In Figure 4D we observe that the I.p. EPG mutant, which adopts the FXF motif, does not have a significant response to the magnetic stimulus which is consistent with previous findings that removing the central F residue (position 7 in the motif) renders the protein non-functional (*15*). The other mutants in Figures 4C, E, & F *do* have the FFF motif, and all exhibit a response to the magnetic stimulus. It is also worth noting that the B.g. EPG mutant displayed the fastest response to the magnetic stimulus compared to the native EPG as can be seen more clearly in Figure 4G – GCaMP6m intensity upticks after pulse number 3 in the B.g. mutant instead of after pulse number 4 like native EPG. Figure 4H demonstrates that mutating the 3F motif of EPG does not disrupt its overall structure. A faster response from EPG is more desirable for certain synthetic applications and for several neuroscience applications (*19, 20*) – and the B.g. mutant shows it is possible to mutate the 3F motif to achieve this.

## Conclusion

Ultimately, this research brings to light more information about EPG. By identifying homologous proteins, we revealed the conservation of a 3F motif among Teleosts. The information presented regarding the potential origins of EPG, and characteristics of fish that may or may not exhibit magnetoreception has significant potential to be expanded upon as genomic and transcriptomic data become available for new species (among Teleostei and other classes). We have also continued to push EPG’s potential as a synthetic tool by identifying a mutation of the 3F motif, which enhances functionality in terms of calcium signaling. This information also encourages further expansion by engineering EPG to function as a control mechanism for specific tasks.

## Methods

### Identifying Homologs via tblastn

The entirety of EPG’s amino acid sequence was used as the query sequence in NCBI’s tblastn. The search was conducted using the Nucleotide collection (nr/nt) database with no additional constraints or modifiers. Amino acid and nucleotide sequences for each unique hit were saved for analysis.

### Amino Acid Alignment

A multiple sequence alignment was conducted in SnapGene version 7.1.2 with EPG and all other identified homologs. Sequences were aligned using MUSCLE under the default settings. The consensus sequence is displayed with a threshold of > 50%. Amino acids matching the consensus were highlighted in green and the sequence conservation was displayed as colored bars.

### Sequence Logo Generation

WebLogo Version 2.8.2 (*22*) was used to generate the sequence logo for the 3F motif of EPG and all homologs. The motifs were pulled from the previously created alignment. A customized color scheme that reflects the Zappo color scheme was used to highlight physical chemical properties within the motif.

### Phylogenetic Tree Construction

Scientific names for each species containing an EPG homolog were uploaded to TimeTree (*23*) in the form of a text file. The output phylogeny was exported with default formatting.

### Obtaining Characteristic Data

Each species with an EPG homolog was searched for in FishBase (*33*) by scientific name. Information regarding the geographical location, water type, migratory habits, depth, water temperature, and maximum length was recorded. Any information not available on FishBase was labeled as undetermined.

### Statistical Analysis of Fish Characteristics

Data regarding migratory habits, water type, depth classification, and water temperature were separated by FFF and FXF motif and displayed in pie charts created using Prism 10. Data regarding max length was displayed as a scatter plot with a median line in Prism 10.

To formally test for the association of the presence/absence of the FFF motif and various species’ morphological or adaptation characteristics we fitted logistic regressions that had the presence/absence of the FFF motif as the response and features of the species as predictors. Separate logistic regressions were fitted for migratory behavior, water type, depth, temperature, and maximum length. To achieve higher power for this analysis, we grouped categories that had a very small number of species into broader categories. All these analyses were done using R (*35*). The logistic regression models were fitted using the glm function. To test the significance of the association of each factor with the presence/absence of the FFF motif we used a Likelihood Ratio test, comparing the fitted models with an intercept only model. For factors that showed significant association (Likelihood Ratio Test p-value < 0.01), we further tested for differences in the frequency of the FFF motif between levels of the factors. For body length, we also tested the association with the presence/absence of the FFF motif using a linear model with body length as the response and the motif as a predictor.

### Plasmid Construction & Site Directed Mutagenesis

A plasmid containing GCaMP6m was obtained from Addgene (*36*) and gblocks were obtained from IDT. The NEBuilder HiFi DNA Assembly kit (NEB) was used to create the plasmid EPG-IRES-GCaMP6m. Site directed mutagenesis of the EPG 3F motif was carried out via PCR with primers obtained from IDT on the EPG-IRES-GCaMP6m plasmid.

### Cell Culture

HeLa cells were cultured with DMEM supplemented with 10% FBS and 1% PenStrep in 25cm^2^ polystyrene flasks and grown in an incubator that maintains humidity, a temperature of 37°C, and 5% CO_2_.

### GCaMP6m Assay

HeLa cells were seeded into 35 mm polystyrene tissue culture dishes at a density of 0.1 × 10^6^ cells. After 24 hours, HeLa cells were transfected with the Lipofectamine 3000 kit according to the manufacturer’s protocol. 24 hours post transfection, HeLa cells were washed twice with 1mL pre-warmed DMEM and covered with 2mL pre-warmed DMEM. HeLa were imaged in the Keyence BZ-X770 microscope equipped with a 10X objective and Tokai kit chamber that maintained the cells at 37°C with 5% CO_2_ and humidity. GCaMP6m was visualized with the Keyence BZ-X GFP filter; the expression of GCaMP6m is also indicative of the expression of the preceding construct (i.e., EPG or an EPG mutant) due to the IRES sequence.

Control groups received no stimulation and remained undisturbed for the 25-minute microscopy duration. Sham groups received four 15-second pulses from a custom air-core coil (*37*) at 4.5A (∼0.3mT) with 5 minutes of rest between each pulse. Active groups received four 15-second pulses from a custom air-core coil at 4.5A (∼14.5mT) with 5 minutes of rest between each pulse. All experiments were conducted in groups of three such that one dish received only one type of stimulus.

### GCaMP6m Assay Data Analysis

Videos of GCaMP6m were split into 50 frames to form a timelapse for analysis. The Time Series Analyzer V3 (*38*) plugin for FIJI (*39*) was used to place ROIs around cells of interest. ROI placement ensured the cell remained in the borders for all 50 frames while minimizing background. Intensity values from FIJI were normalized to the first point in the read to make data more easily interpretable. Data gathered from each ROI over all experiments for a specific construct were averaged and plotted using Prism 10 to display the average intensity of GCaMP6m over time with error bars as 95% CI.

### Structure Prediction

Amino acid sequences were input to Robetta (*40*) and structures were predicted using the default settings. Structures were aligned and visualized using PyMOL (*34*).

## Supporting information

Ricker et al. EPG Homology SM

TimeTree Fish Species

EPG Homologs Table

Fish Homology Condensed

Fish Homology NT

GCaMP Assay Raw Data

## Acknowledgments

A.A.G. acknowledges financial support from the NIH/NINDS: R01-NS098231; R01-NS104306 and NIH/NIBIB: R01-EB031008; R01-EB030565; R01-EB031936.

## Notes

### Competing Interest Statement

The authors have declared no competing interest.

### Summary of Updates

The author list has been updated.

