## Supplementary material for "A conserved phenylalanine motif among Teleost fish provides insight for improving electromagnetic perception": Ricker et al. EPG Homology SM

### Supplemental Material

In addition to the supplemental files, please find below some figures to support our findings. We have also provided below the DNA sequences for the constructs used in this project.

#### Supplemental Files:

*TimeTree Fish Species*: A text file containing species names that was used to generate the phylogenetic tree shown in Figure 2.

*EPG Homologs Table*: A spreadsheet organizing all the information gathered about species with an EPG homolog throughout the duration of the project.

*Fish Homology Condensed*: A spreadsheet condensing some information from EPG Homologs Table which was used to conduct the statistical analysis for Figures S1-S4 and Tables S1-S4.

*Fish Homology NT*: A spreadsheet detailing the nucleotide sequences of the EPG homologs.

*GCaMP Assay Raw Data*: A spreadsheet containing raw data from all the GCaMP6m experiments displayed in Figure 4 and Figure S5.

#### Supplemental Figures:

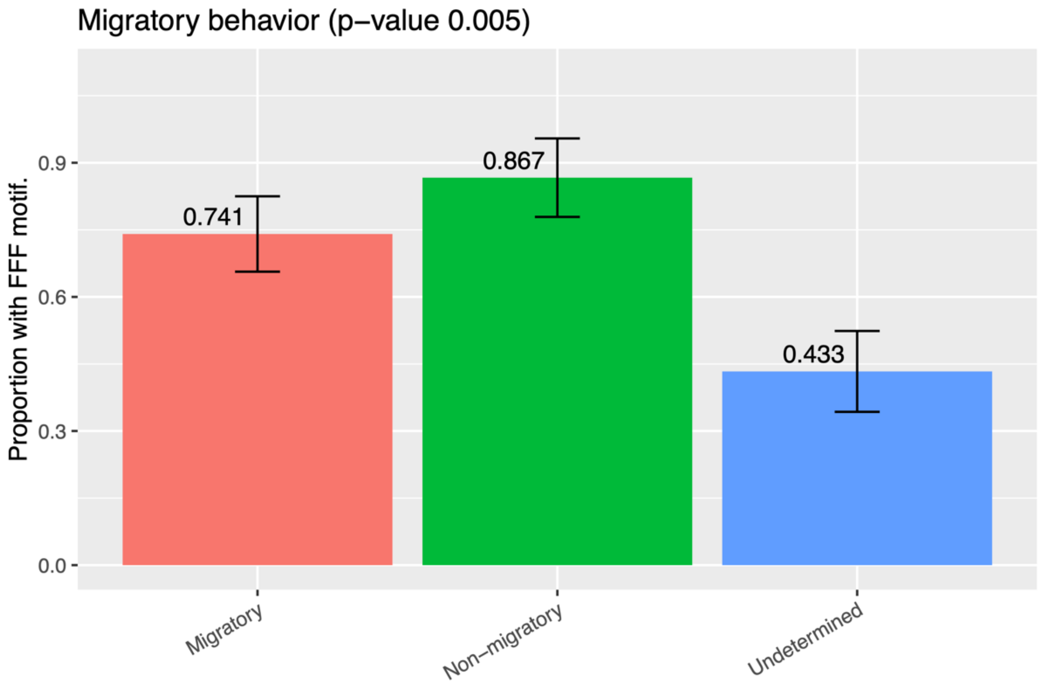


**Figure S1. Frequency (+/- SE) of the FFF motif by migratory behavior.** Displayed in conjunction with Figure 3A. The migratory group includes oceanodromous, anadromous, amphidromous, catadromous, and potadromous species.

**Table S1: Differences in the frequency of the FFF motif between groups (below the diagonal) and p-value for the difference (above the diagonal).**

| | Migratory | Non-migratory | Undetermined |
|---|---|---|---|
| **Migratory** | – | 0.34883 | 0.02148 |
| **Non-migratory** | -0.12593 | – | 0.01124 |
| **Undetermined** | 0.30741 | 0.43333 | – |

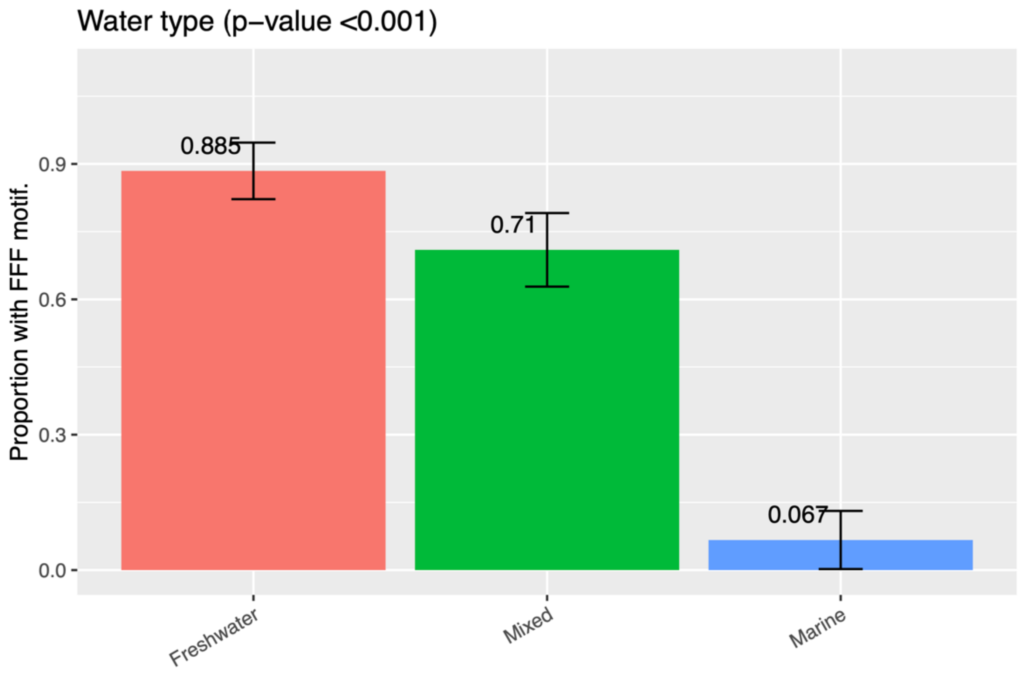


**Figure S2. Frequency (+/- SE) of the FFF motif by water type.** Displayed in conjunction with Figure 3B. The mixed group includes species that exist in multiple water types or have brackish tolerance.

**Table S2: Differences in the frequency of the FFF motif between groups (below the diagonal) and p-value for the difference (above the diagonal).**

| | Freshwater | Mixed | Marine |
|---|---|---|---|
| **Freshwater** | – | 0.11755 | 1e-04 |
| **Mixed** | 0.17494 | – | 0.00143 |
| **Marine** | 0.81795 | 0.64301 | – |

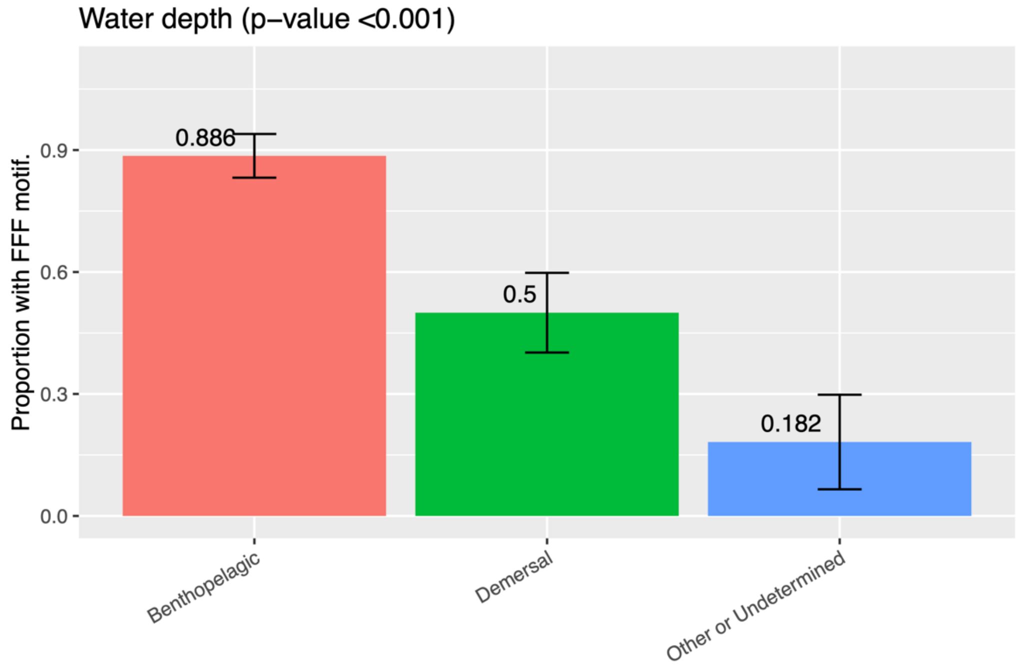


**Figure S3. Frequency (+/- SE) of the FFF motif by water depth.** Displayed in conjunction with Figure 3C. The other and undetermined group contains species that exist in pelagic or bathydemersal zones and species that are undetermined.

**Table S3: Differences in the frequency of the FFF motif between groups (below the diagonal) and p-value for the difference (above the diagonal).**

| | Benthopelagic | Demersal | Other or Undetermined |
|---|---|---|---|
| **Benthopelagic** | – | 0.00193 | 0.00017 |
| **Demersal** | 0.38571 | – | 0.08549 |
| **Other or Undetermined** | 0.7039 | 0.31818 | – |

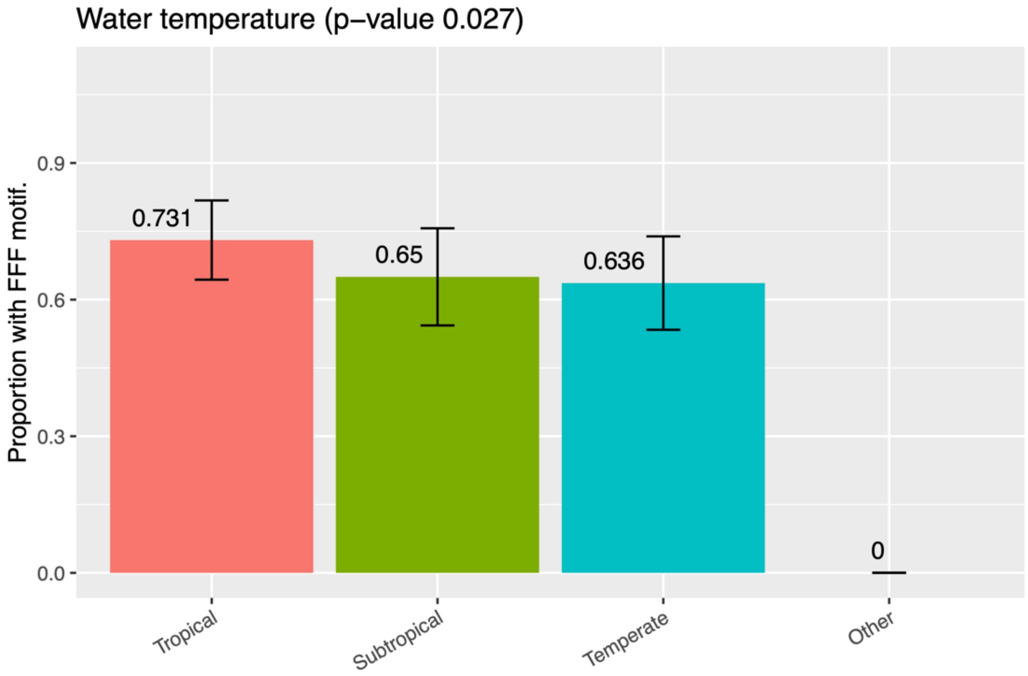


**Figure S4. Frequency (+/- SE) of the FFF motif by water temperature.** Displayed in conjunction with Figure 3D. The other group encompasses polar and deep-water designations.

**Table S4: Parameter estimates, SE, and p-values for the logistic regression of the presence/absence of the FFF motif on maximum body length.**

| | Estimate | Std. Error | z value | Pr(>\|z\|) |
|---|---|---|---|---|
| **(Intercept)** | 0.6062 | 0.3631 | 1.6697 | 0.0950 |
| **Max Body Length** | -0.0005 | 0.0034 | -0.1339 | 0.8935 |

**Table S5: Parameter estimates, SE, and p-values for the linear regression of maximum recorded length on the presence/absence of the FFF motif.**

| | Estimate | Std. Error | t value | Pr(>\|t\|) |
|---|---|---|---|---|
| **(Intercept)** | 78.831 | 14.156 | 5.569 | 0.000 |
| **Motif FFF** | -2.339 | 17.710 | -0.132 | 0.895 |

**Table S6: Percentage of fish shorter and longer than 20cm by motif type.**

| | <20cm | >/=20cm |
|---|---|---|
| **FXF** | 19.2 | 80.8 |
| **FFF** | 43.5 | 56.5 |

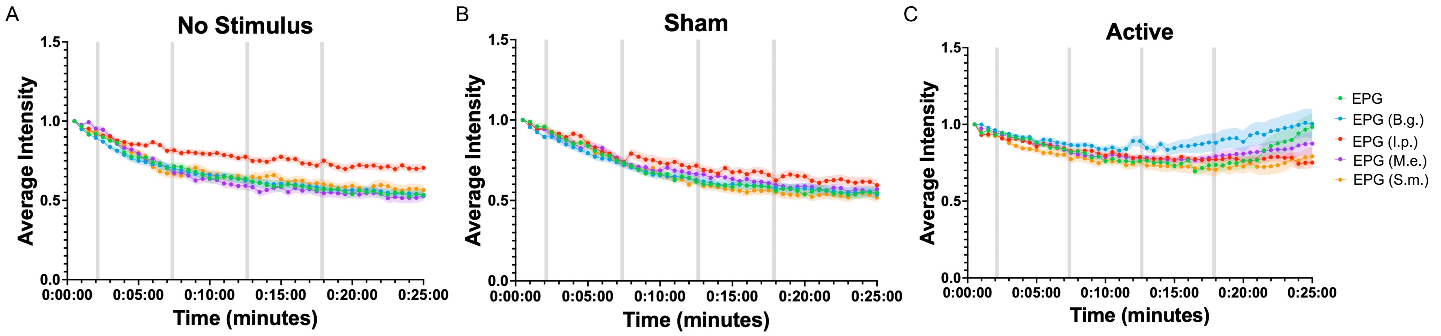


**Figure S5. GCaMP6m assay with EPG and EPG 3F mutants based on homologs.** (A-C) This is the same data presented in Figure 4, grouped by stimulus rather than construct. HeLa cells expressing versions of EPG-IRES-GCaMP6m subjected to a GCaMP6m functional assay; cells were exposed to either active, sham, or no stimulus with a pulse pattern of 15s on 5min off for 4 pulses (gray bars). Error bars are representative of 95% CI. Experiments include n=90 cells over three experiments for each no stimulus, sham, and active groups respectively. (A) No stimulus groups. (B) Sham groups. (C) Active groups.

### Plasmid DNA Sequences:

#### EPG-IRES-GCaMP6m:

GACGGATCGGGAGATCTCCCGATCCCCTATGGTCGACTCTCAGTACAATCTGCTCTGATGCCGCATAGTTAAGCCAGTATCTGCTCCCTGCTTGTGTGTTGGAGGTCGCTGA
GTAGTGCGCGAGCAAAATTTAAGCTACAACAAGGCAAGGCTTGACCGACAATTGCATGAAGAATCTGCTTAGGGTTAGGCGTTTTGCGCTGCTTCGCGATGTACGGGCCAGA
TATACGCGTTGACATTGATTATTGACTAGTTATTAATAGTAATCAATTACGGGGTCATTAGTTCATAGCCCATATATGGAGTTCCGCGTTACATAACTTACGGTAAATGGCC
CGCCTGGCTGACCGCCCAACGACCCCCGCCCATTGACGTCAATAATGACGTATGTTCCCATAGTAACGCCAATAGGGACTTTCCATTGACGTCAATGGGTGGACTATTTACG
GTAAACTGCCCACTTGGCAGTACATCAAGTGTATCATATGCCAAGTACGCCCCCTATTGACGTCAATGACGGTAAATGGCCCGCCTGGCATTATGCCCAGTACATGACCTTA
TGGGACTTTCCTACTTGGCAGTACATCTACGTATTAGTCATCGCTATTACCATGGTGATGCGGTTTTGGCAGTACATCAATGGGCGTGGATAGCGGTTTGACTCACGGGGAT
TTCCAAGTCTCCACCCCATTGACGTCAATGGGAGTTTGTTTTGGCACCAAAATCAACGGGACTTTCCAAAATGTCGTAACAACTCCGCCCCATTGACGCAAATGGGCGGTAG
GCGTGTACGGTGGGAGGTCTATATAAGCAGAGCTCTCTGGCTAACTAGAGAACCCACTGCTTACTGGCTTATCGAAATTAATACGACTCACTATAGGGAGACCCAAGCTGGC
TAGTTAAGCTTGGTACCGAGCTCGGATCCACCATGAAGTGTGTACTTTTGGGATTCGCAGCAGTGATCGGATTCTTCGCGATCGCGGAGTCTCTTACCTGTAACACATGCTC
AGTGAGTCTGATTGGAATATGTCTGAATCCCGCAACAGCGACTTGCTCCACCAACACATCCGTCTGCACCACAGGAAGAGCCAGTTTCACGGGCGTCCTCGGCTTCCTGGGC
TTCAACTCCCAGGGCTGCACGGAGGGAGCTCAGTGTAATGGCACCGTGTCCGGGTCCATCCTGGGTGCGTCGTACACGGTCACTCAAACCTGCTGCAGCACAAACAACTGCA
ACCCCGTGACCAGCGGCGCCTCCTACGTCCAGATCTCCGTCAGCGCGGCCCTGAGCGCCGCCCTGCTGGCCTGCGTCTGGGGCCAGTCCGTCTACGACTACAAGGATGACGA
TGACAAGGATTACAAAGACGACGATGATAAGGACTATAAGGATGATGACGACAAATAATAGCAATTCCTCGACGACTGCATAGGGTTACCCCCCTCTCCCTCCCCCCCCCCT
AACGTTACTGGCCGAAGCCGCTTGGAATAAGGCCGGTGTGCGTTTGTCTATATGTTATTTTCCACCATATTGCCGTCTTTTGGCAATGTGAGGGCCCGGAAACCTGGCCCTG
TCTTCTTGACGAGCATTCCTAGGGGTCTTTCCCCTCTCGCCAAAGGAATGCAAGGTCTGTTGAATGTCGTGAAGGAAGCAGTTCCTCTGGAAGCTTCTTGAAGACAAACAAC
GTCTGTAGCGACCCTTTGCAGGCAGCGGAACCCCCCACCTGGCGACAGGTGCCTCTGCGGCCAAAAGCCACGTGTATAAGATACACCTGCAAAGGCGGCACAACCCCAGTGC
CACGTTGTGAGTTGGATAGTTGTGGAAAGAGTCAAATGGCTCTCCTCAAGCGTATTCAACAAGGGGCTGAAGGATGCCCAGAAGGTACCCCATTGTATGGGATCTGATCTGG
GGCCTCGGTGCACATGCTTTACATGTGTTTAGTCGAGGTTAAAAAACGTCTAGGCCCCCCGAACCACGGGGACGTGGTTTTCCTTTGAAAAACACGATGATAATGGCCACAA
CCATGGGTTCTCATCATCATCATCATCATGGTATGGCTAGCATGACTGGTGGACAGCAAATGGGTCGGGATCTGTACGACGATGACGATAAGGATCTCGCCACCATGGTCGA
CTCATCACGTCGTAAGTGGAATAAGACAGGTCACGCAGTCAGAGCTATAGGTCGGCTGAGCTCACTCGAGAACGTCTATATCAAGGCCGACAAGCAGAAGAACGGCATCAAG
GCGAACTTCAAGATCCGCCACAACATCGAGGACGGCGGCGTGCAGCTCGCCTACCACTACCAGCAGAACACCCCCATCGGCGACGGCCCCGTGCTGCTGCCCGACAACCACT
ACCTGAGCGTGCAGTCCAAACTTTCGAAAGACCCCAACGAGAAGCGCGATCACATGGTCCTGCTGGAGTTCGTGACCGCCGCCGGGATCACTCTCGGCATGGACGAGCTGTA
CAAGGGCGGTACCGGAGGGAGCATGGTGAGCAAGGGCGAGGAGCTGTTCACCGGGGTGGTGCCCATCCTGGTCGAGCTGGACGGCGACGTAAACGGCCACAAGTTCAGCGTG
TCCGGCGAGGGTGAGGGCGATGCCACCTACGGCAAGCTGACCCTGAAGTTCATCTGCACCACCGGCAAGCTGCCCGTGCCCTGGCCCACCCTCGTGACCACCCTGACCTACG
GCGTGCAGTGCTTCAGCCGCTACCCCGACCACATGAAGCAGCACGACTTCTTCAAGTCCGCCATGCCCGAAGGCTACATCCAGGAGCGCACCATCTTCTTCAAGGACGACGG
CAACTACAAGACCCGCGCCGAGGTGAAGTTCGAGGGCGACACCCTGGTGAACCGCATCGAGCTGAAGGGCATCGACTTCAAGGAGGACGGCAACATCCTGGGGCACAAGCTG
GAGTACAACCTGCCGGACCAACTGACTGAAGAGCAGATCGCAGAATTTAAAGAGGCTTTCTCCCTATTTGACAAGGACGGGGATGGGACAATAACAACCAAGGAGCTGGGGA
CGGTGATGCGGTCTCTGGGGCAGAACCCCACAGAAGCAGAGCTGCAGGACATGATCAATGAAGTAGATGCCGACGGTGACGGCACAATCGACTTCCCTGAGTTCCTGACAAT
GATGGCAAGAAAAGGGAGCTACAGGGACACGGAAGAAGAAATTAGAGAAGCGTTCGGTGTGTTTGATAAGGATGGCAATGGCTACATCAGTGCAGCAGAGCTTCGCCACGTG
ATGACAAACCTTGGAGAGAAGTTAACAGATGAAGAGGTTGATGAAATGATCAGGGAAGCAGACATCGATGGGGATGGTCAGGTAAACTACGAAGAGTTTGTACAAATGATGA
CAGCGAAGTAATCTAGAGGGCCCTTCGAAGGTAAGCCTATCCCTAACCCTCTCCTCGGTCTCGATTCTACGCGTACCGGTCATCATCACCATCACCATTGAGTTTAAACCCG
CTGATCAGCCTCGACTGTGCCTTCTAGTTGCCAGCCATCTGTTGTTTGCCCCTCCCCCGTGCCTTCCTTGACCCTGGAAGGTGCCACTCCCACTGTCCTTTCCTAATAAAAT
GAGGAAATTGCATCGCATTGTCTGAGTAGGTGTCATTCTATTCTGGGGGGTGGGGTGGGGCAGGACAGCAAGGGGGAGGATTGGGAAGACAATAGCAGGCATGCTGGGGATG
CGGTGGGCTCTATGGCTTCTGAGGCGGAAAGAACCAGCTGGGGCTCTAGGGGGTATCCCCACGCGCCCTGTAGCGGCGCATTAAGCGCGGCGGGTGTGGTGGTTACGCGCAG
CGTGACCGCTACACTTGCCAGCGCCCTAGCGCCCGCTCCTTTCGCTTTCTTCCCTTCCTTTCTCGCCACGTTCGCCGGCTTTCCCCGTCAAGCTCTAAATCGGGGCATCCCT
TTAGGGTTCCGATTTAGTGCTTTACGGCACCTCGACCCCAAAAAACTTGATTAGGGTGATGGTTCACGTAGTGGGCCATCGCCCTGATAGACGGTTTTTCGCCCTTTGACGT
TGGAGTCCACGTTCTTTAATAGTGGACTCTTGTTCCAAACTGGAACAACACTCAACCCTATCTCGGTCTATTCTTTTGATTTATAAGGGATTTTGGGGATTTCGGCCTATTG
GTTAAAAAATGAGCTGATTTAACAAAAATTTAACGCGAATTAATTCTGTGGAATGTGTGTCAGTTAGGGTGTGGAAAGTCCCCAGGCTCCCCAGGCAGGCAGAAGTATGCAA
AGCATGCATCTCAATTAGTCAGCAACCAGGTGTGGAAAGTCCCCAGGCTCCCCAGCAGGCAGAAGTATGCAAAGCATGCATCTCAATTAGTCAGCAACCATAGTCCCGCCCC
TAACTCCGCCCATCCCGCCCCTAACTCCGCCCAGTTCCGCCCATTCTCCGCCCCATGGCTGACTAATTTTTTTTATTTATGCAGAGGCCGAGGCCGCCTCTGCCTCTGAGCT
ATTCCAGAAGTAGTGAGGAGGCTTTTTTGGAGGCCTAGGCTTTTGCAAAAAGCTCCCGGGAGCTTGTATATCCATTTTCGGATCTGATCAAGAGACAGGATGAGGATCGTTT
CGCATGATTGAACAAGATGGATTGCACGCAGGTTCTCCGGCCGCTTGGGTGGAGAGGCTATTCGGCTATGACTGGGCACAACAGACAATCGGCTGCTCTGATGCCGCCGTGT
TCCGGCTGTCAGCGCAGGGGCGCCCGGTTCTTTTTGTCAAGACCGACCTGTCCGGTGCCCTGAATGAACTGCAGGACGAGGCAGCGCGGCTATCGTGGCTGGCCACGACGGG
CGTTCCTTGCGCAGCTGTGCTCGACGTTGTCACTGAAGCGGGAAGGGACTGGCTGCTATTGGGCGAAGTGCCGGGGCAGGATCTCCTGTCATCTCACCTTGCTCCTGCCGAG
AAAGTATCCATCATGGCTGATGCAATGCGGCGGCTGCATACGCTTGATCCGGCTACCTGCCCATTCGACCACCAAGCGAAACATCGCATCGAGCGAGCACGTACTCGGATGG
AAGCCGGTCTTGTCGATCAGGATGATCTGGACGAAGAGCATCAGGGGCTCGCGCCAGCCGAACTGTTCGCCAGGCTCAAGGCGCGCATGCCCGACGGCGAGGATCTCGTCGT
GACCCATGGCGATGCCTGCTTGCCGAATATCATGGTGGAAAATGGCCGCTTTTCTGGATTCATCGACTGTGGCCGGCTGGGTGTGGCGGACCGCTATCAGGACATAGCGTTG

GCTACCCGTGATATTGCTGAAGAGCTTGGCGGCGAATGGGCTGACCGCTTCCTCGTGCTTTACGGTATCGCCGCTCCCGATTCGCAGCGCATCGCCTTCTATCGCCTTCTTG
ACGAGTTCTTCTGAGCGGGACTCTGGGGTTCGCGAAATGACCGACCAAGCGACGCCCAACCTGCCATCACGAGATTTCGATTCCACCGCCGCCTTCTATGAAAGGTTGGGCT
TCGGAATCGTTTTCCGGGACGCCGGCTGGATGATCCTCCAGCGCGGGGATCTCATGCTGGAGTTCTTCGCCCACCCCAACTTGTTTATTGCAGCTTATAATGGTTACAAATA
AAGCAATAGCATCACAAATTTCACAAATAAAGCATTTTTTTCACTGCATTCTAGTTGTGGTTTGTCCAAACTCATCAATGTATCTTATCATGTCTGTATACCGTCGACCTCT
AGCTAGAGCTTGGCGTAATCATGGTCATAGCTGTTTCCTGTGTGAAATTGTTATCCGCTCACAATTCCACACAACATACGAGCCGGAAGCATAAAGTGTAAAGCCTGGGGTG
CCTAATGAGTGAGCTAACTCACATTAATTGCGTTGCGCTCACTGCCCGCTTTCCAGTCGGGAAACCTGTCGTGCCAGCTGCATTAATGAATCGGCCAACGCGCGGGGAGAGG
CGGTTTGCGTATTGGGCGCTCTTCCGCTTCCTCGCTCACTGACTCGCTGCGCTCGGTCGTTCGGCTGCGGCGAGCGGTATCAGCTCACTCAAAGGCGGTAATACGGTTATCC
ACAGAATCAGGGGATAACGCAGGAAAGAACATGTGAGCAAAAGGCCAGCAAAAGGCCAGGAACCGTAAAAAGGCCGCGTTGCTGGCGTTTTTCCATAGGCTCCGCCCCCCTG
ACGAGCATCACAAAAATCGACGCTCAAGTCAGAGGTGGCGAAACCCGACAGGACTATAAAGATACCAGGCGTTTCCCCCTGGAAGCTCCCTCGTGCGCTCTCCTGTTCCGAC
CCTGCCGCTTACCGGATACCTGTCCGCCTTTCTCCCTTCGGGAAGCGTGGCGCTTTCTCAATGCTCACGCTGTAGGTATCTCAGTTCGGTGTAGGTCGTTCGCTCCAAGCTG
GGCTGTGTGCACGAACCCCCCGTTCAGCCCGACCGCTGCGCCTTATCCGGTAACTATCGTCTTGAGTCCAACCCGGTAAGACACGACTTATCGCCACTGGCAGCAGCCACTG
GTAACAGGATTAGCAGAGCGAGGTATGTAGGCGGTGCTACAGAGTTCTTGAAGTGGTGGCCTAACTACGGCTACACTAGAAGGACAGTATTTGGTATCTGCGCTCTGCTGAA
GCCAGTTACCTTCGGAAAAAGAGTTGGTAGCTCTTGATCCGGCAAACAAACCACCGCTGGTAGCGGTGGTTTTTTTGTTTGCAAGCAGCAGATTACGCGCAGAAAAAAAGGA
TCTCAAGAAGATCCTTTGATCTTTTCTACGGGGTCTGACGCTCAGTGGAACGAAAACTCACGTTAAGGGATTTTGGTCATGAGATTATCAAAAAGGATCTTCACCTAGATCC
TTTTAAATTAAAAATGAAGTTTTAAATCAATCTAAAGTATATATGAGTAAACTTGGTCTGACAGTTACCAATGCTTAATCAGTGAGGCACCTATCTCAGCGATCTGTCTATT
TCGTTCATCCATAGTTGCCTGACTCCCCGTCGTGTAGATAACTACGATACGGGAGGGCTTACCATCTGGCCCCAGTGCTGCAATGATACCGCGAGACCCACGCTCACCGGCT
CCAGATTTATCAGCAATAAACCAGCCAGCCGGAAGGGCCGAGCGCAGAAGTGGTCCTGCAACTTTATCCGCCTCCATCCAGTCTATTAATTGTTGCCGGGAAGCTAGAGTAA
GTAGTTCGCCAGTTAATAGTTTGCGCAACGTTGTTGCCATTGCTACAGGCATCGTGGTGTCACGCTCGTCGTTTGGTATGGCTTCATTCAGCTCCGGTTCCCAACGATCAAG
GCGAGTTACATGATCCCCCATGTTGTGCAAAAAAGCGGTTAGCTCCTTCGGTCCTCCGATCGTTGTCAGAAGTAAGTTGGCCGCAGTGTTATCACTCATGGTTATGGCAGCA
CTGCATAATTCTCTTACTGTCATGCCATCCGTAAGATGCTTTTCTGTGACTGGTGAGTACTCAACCAAGTCATTCTGAGAATAGTGTATGCGGCGACCGAGTTGCTCTTGCC
CGGCGTCAATACGGGATAATACCGCGCCACATAGCAGAACTTTAAAAGTGCTCATCATTGGAAAACGTTCTTCGGGGCGAAAACTCTCAAGGATCTTACCGCTGTTGAGATC
CAGTTCGATGTAACCCACTCGTGCACCCAACTGATCTTCAGCATCTTTTACTTTCACCAGCGTTTCTGGGTGAGCAAAAACAGGAAGGCAAAATGCCGCAAAAAAGGGAATA
AGGGCGACACGGAAATGTTGAATACTCATACTCTTCCTTTTTCAATATTATTGAAGCATTTATCAGGGTTATTGTCTCATGAGCGGATACATATTTGAATGTATTTAGAAAA
ATAAACAAATAGGGGTTCCGCGCACATTTCCCCGAAAAGTGCCACCTGACGTC

#### EPG (B.g. 3F)-IRES-GCaMP6m:

GACGGATCGGGAGATCTCCCGATCCCCTATGGTCGACTCTCAGTACAATCTGCTCTGATGCCGCATAGTTAAGCCAGTATCTGCTCCCTGCTTGTGTGTTGGAGGTCGCTGA
GTAGTGCGCGAGCAAAATTTAAGCTACAACAAGGCAAGGCTTGACCGACAATTGCATGAAGAATCTGCTTAGGGTTAGGCGTTTTGCGCTGCTTCGCGATGTACGGGCCAGA
TATACGCGTTGACATTGATTATTGACTAGTTATTAATAGTAATCAATTACGGGGTCATTAGTTCATAGCCCATATATGGAGTTCCGCGTTACATAACTTACGGTAAATGGCC
CGCCTGGCTGACCGCCCAACGACCCCCGCCCATTGACGTCAATAATGACGTATGTTCCCATAGTAACGCCAATAGGGACTTTCCATTGACGTCAATGGGTGGACTATTTACG
GTAAACTGCCCACTTGGCAGTACATCAAGTGTATCATATGCCAAGTACGCCCCCTATTGACGTCAATGACGGTAAATGGCCCGCCTGGCATTATGCCCAGTACATGACCTTA
TGGGACTTTCCTACTTGGCAGTACATCTACGTATTAGTCATCGCTATTACCATGGTGATGCGGTTTTGGCAGTACATCAATGGGCGTGGATAGCGGTTTGACTCACGGGGAT
TTCCAAGTCTCCACCCCATTGACGTCAATGGGAGTTTGTTTTGGCACCAAAATCAACGGGACTTTCCAAAATGTCGTAACAACTCCGCCCCATTGACGCAAATGGGCGGTAG
GCGTGTACGGTGGGAGGTCTATATAAGCAGAGCTCTCTGGCTAACTAGAGAACCCACTGCTTACTGGCTTATCGAAATTAATACGACTCACTATAGGGAGACCCAAGCTGGC
TAGTTAAGCTTGGTACCGAGCTCGGATCCACTAGTCCAGTGTGGTGGAATTGCCCTTCCAGTACCCTTCACCATGAAGTGTGTACTTTTGGGATTCGCAGCAGTGATCGGAT
TCTTCGCGATCGCGGAGTCTCTTACCTGTAACACATGCTCAGTGAGTCTGATTGGAATATGTCTGAATCCCGCAACAGCGACTTGCTCCACCAACACATCCGTCTGCACCAC
AGGAAGAGCCAGTTTCCCGTCTATCTCTGGTTTCCTGGGTTTCAACTCCCAGGGCTGCACGGAGGGAGCTCAGTGTAATGGCACCGTGTCCGGGTCCATCCTGGGTGCGTCG
TACACGGTCACTCAAACCTGCTGCAGCACAAACAACTGCAACCCCGTGACCAGCGGCGCCTCCTACGTCCAGATCTCCGTCAGCGCGGCCCTGAGCGCCGCCCTGCTGGCCT
GCGTCTGGGGCCAGTCCGTCTACGACTACAAGGATGACGATGACAAGGATTACAAAGACGACGATGATAAGGACTATAAGGATGATGACGACAAATAATAGCAATTCCTCGA
CGACTGCATAGGGTTACCCCCCTCTCCCTCCCCCCCCCCTAACGTTACTGGCCGAAGCCGCTTGGAATAAGGCCGGTGTGCGTTTGTCTATATGTTATTTTCCACCATATTG
CCGTCTTTTGGCAATGTGAGGGCCCGGAAACCTGGCCCTGTCTTCTTGACGAGCATTCCTAGGGGTCTTTCCCCTCTCGCCAAAGGAATGCAAGGTCTGTTGAATGTCGTGA
AGGAAGCAGTTCCTCTGGAAGCTTCTTGAAGACAAACAACGTCTGTAGCGACCCTTTGCAGGCAGCGGAACCCCCCACCTGGCGACAGGTGCCTCTGCGGCCAAAAGCCACG
TGTATAAGATACACCTGCAAAGGCGGCACAACCCCAGTGCCACGTTGTGAGTTGGATAGTTGTGGAAAGAGTCAAATGGCTCTCCTCAAGCGTATTCAACAAGGGGCTGAAG
GATGCCCAGAAGGTACCCCATTGTATGGGATCTGATCTGGGGCCTCGGTGCACATGCTTTACATGTGTTTAGTCGAGGTTAAAAAACGTCTAGGCCCCCCGAACCACGGGGA
CGTGGTTTTCCTTTGAAAAACACGATGATAATGGCCACAACCATGGGTTCTCATCATCATCATCATCATGGTATGGCTAGCATGACTGGTGGACAGCAAATGGGTCGGGATC
TGTACGACGATGACGATAAGGATCTCGCCACCATGGTCGACTCATCACGTCGTAAGTGGAATAAGACAGGTCACGCAGTCAGAGCTATAGGTCGGCTGAGCTCACTCGAGAA
CGTCTATATCAAGGCCGACAAGCAGAAGAACGGCATCAAGGCGAACTTCAAGATCCGCCACAACATCGAGGACGGCGGCGTGCAGCTCGCCTACCACTACCAGCAGAACACC
CCCATCGGCGACGGCCCCGTGCTGCTGCCCGACAACCACTACCTGAGCGTGCAGTCCAAACTTTCGAAAGACCCCAACGAGAAGCGCGATCACATGGTCCTGCTGGAGTTCG
TGACCGCCGCCGGGATCACTCTCGGCATGGACGAGCTGTACAAGGGCGGTACCGGAGGGAGCATGGTGAGCAAGGGCGAGGAGCTGTTCACCGGGGTGGTGCCCATCCTGGT
CGAGCTGGACGGCGACGTAAACGGCCACAAGTTCAGCGTGTCCGGCGAGGGTGAGGGCGATGCCACCTACGGCAAGCTGACCCTGAAGTTCATCTGCACCACCGGCAAGCTG
CCCGTGCCCTGGCCCACCCTCGTGACCACCCTGACCTACGGCGTGCAGTGCTTCAGCCGCTACCCCGACCACATGAAGCAGCACGACTTCTTCAAGTCCGCCATGCCCGAAG
GCTACATCCAGGAGCGCACCATCTTCTTCAAGGACGACGGCAACTACAAGACCCGCGCCGAGGTGAAGTTCGAGGGCGACACCCTGGTGAACCGCATCGAGCTGAAGGGCAT
CGACTTCAAGGAGGACGGCAACATCCTGGGGCACAAGCTGGAGTACAACCTGCCGGACCAACTGACTGAAGAGCAGATCGCAGAATTTAAAGAGGCTTTCTCCCTATTTGAC
AAGGACGGGGATGGGACAATAACAACCAAGGAGCTGGGGACGGTGATGCGGTCTCTGGGGCAGAACCCCACAGAAGCAGAGCTGCAGGACATGATCAATGAAGTAGATGCCG
ACGGTGACGGCACAATCGACTTCCCTGAGTTCCTGACAATGATGGCAAGAAAAGGGAGCTACAGGGACACGGAAGAAGAAATTAGAGAAGCGTTCGGTGTGTTTGATAAGGA
TGGCAATGGCTACATCAGTGCAGCAGAGCTTCGCCACGTGATGACAAACCTTGGAGAGAAGTTAACAGATGAAGAGGTTGATGAAATGATCAGGGAAGCAGACATCGATGGG
GATGGTCAGGTAAACTACGAAGAGTTTGTACAAATGATGACAGCGAAGTGAGAATTCTGCAGATATAAGGGCAATTCTGCAGATATCCAGCACAGTGGCGGCCGCTCGAGTC
TAGAGGGCCCGCGGTTCGAAGGTAAGCCTATCCCTAACCCTCTCCTCGGTCTCGATTCTACGCGTACCGGTCATCATCACCATCACCATTGAGTTTAAACCCGCTGATCAGC
CTCGACTGTGCCTTCTAGTTGCCAGCCATCTGTTGTTTGCCCCTCCCCCGTGCCTTCCTTGACCCTGGAAGGTGCCACTCCCACTGTCCTTTCCTAATAAAATGAGGAAATT
GCATCGCATTGTCTGAGTAGGTGTCATTCTATTCTGGGGGGTGGGGTGGGGCAGGACAGCAAGGGGGAGGATTGGGAAGACAATAGCAGGCATGCTGGGGATGCGGTGGGCT
CTATGGCTTCTGAGGCGGAAAGAACCAGCTGGGGCTCTAGGGGGTATCCCCACGCGCCCTGTAGCGGCGCATTAAGCGCGGCGGGTGTGGTGGTTACGCGCAGCGTGACCGC
TACACTTGCCAGCGCCCTAGCGCCCGCTCCTTTCGCTTTCTTCCCTTCCTTTCTCGCCACGTTCGCCGGCTTTCCCCGTCAAGCTCTAAATCGGGGCATCCCTTTAGGGTTC
CGATTTAGTGCTTTACGGCACCTCGACCCCAAAAAACTTGATTAGGGTGATGGTTCACGTAGTGGGCCATCGCCCTGATAGACGGTTTTTCGCCCTTTGACGTTGGAGTCCA
CGTTCTTTAATAGTGGACTCTTGTTCCAAACTGGAACAACACTCAACCCTATCTCGGTCTATTCTTTTGATTTATAAGGGATTTTGGGGATTTCGGCCTATTGGTTAAAAAA
TGAGCTGATTTAACAAAAATTTAACGCGAATTAATTCTGTGGAATGTGTGTCAGTTAGGGTGTGGAAAGTCCCCAGGCTCCCCAGGCAGGCAGAAGTATGCAAAGCATGCAT
CTCAATTAGTCAGCAACCAGGTGTGGAAAGTCCCCAGGCTCCCCAGCAGGCAGAAGTATGCAAAGCATGCATCTCAATTAGTCAGCAACCATAGTCCCGCCCCTAACTCCGC
CCATCCCGCCCCTAACTCCGCCCAGTTCCGCCCATTCTCCGCCCCATGGCTGACTAATTTTTTTTATTTATGCAGAGGCCGAGGCCGCCTCTGCCTCTGAGCTATTCCAGAA
GTAGTGAGGAGGCTTTTTTGGAGGCCTAGGCTTTTGCAAAAAGCTCCCGGGAGCTTGTATATCCATTTTCGGATCTGATCAAGAGACAGGATGAGGATCGTTTCGCATGATT
GAACAAGATGGATTGCACGCAGGTTCTCCGGCCGCTTGGGTGGAGAGGCTATTCGGCTATGACTGGGCACAACAGACAATCGGCTGCTCTGATGCCGCCGTGTTCCGGCTGT
CAGCGCAGGGGCGCCCGGTTCTTTTTGTCAAGACCGACCTGTCCGGTGCCCTGAATGAACTGCAGGACGAGGCAGCGCGGCTATCGTGGCTGGCCACGACGGGCGTTCCTTG
CGCAGCTGTGCTCGACGTTGTCACTGAAGCGGGAAGGGACTGGCTGCTATTGGGCGAAGTGCCGGGGCAGGATCTCCTGTCATCTCACCTTGCTCCTGCCGAGAAAGTATCC
ATCATGGCTGATGCAATGCGGCGGCTGCATACGCTTGATCCGGCTACCTGCCCATTCGACCACCAAGCGAAACATCGCATCGAGCGAGCACGTACTCGGATGGAAGCCGGTC
TTGTCGATCAGGATGATCTGGACGAAGAGCATCAGGGGCTCGCGCCAGCCGAACTGTTCGCCAGGCTCAAGGCGCGCATGCCCGACGGCGAGGATCTCGTCGTGACCCATGG
CGATGCCTGCTTGCCGAATATCATGGTGGAAAATGGCCGCTTTTCTGGATTCATCGACTGTGGCCGGCTGGGTGTGGCGGACCGCTATCAGGACATAGCGTTGGCTACCCGT
GATATTGCTGAAGAGCTTGGCGGCGAATGGGCTGACCGCTTCCTCGTGCTTTACGGTATCGCCGCTCCCGATTCGCAGCGCATCGCCTTCTATCGCCTTCTTGACGAGTTCT
TCTGAGCGGGACTCTGGGGTTCGCGAAATGACCGACCAAGCGACGCCCAACCTGCCATCACGAGATTTCGATTCCACCGCCGCCTTCTATGAAAGGTTGGGCTTCGGAATCG
TTTTCCGGGACGCCGGCTGGATGATCCTCCAGCGCGGGGATCTCATGCTGGAGTTCTTCGCCCACCCCAACTTGTTTATTGCAGCTTATAATGGTTACAAATAAAGCAATAG
CATCACAAATTTCACAAATAAAGCATTTTTTTCACTGCATTCTAGTTGTGGTTTGTCCAAACTCATCAATGTATCTTATCATGTCTGTATACCGTCGACCTCTAGCTAGAGC
TTGGCGTAATCATGGTCATAGCTGTTTCCTGTGTGAAATTGTTATCCGCTCACAATTCCACACAACATACGAGCCGGAAGCATAAAGTGTAAAGCCTGGGGTGCCTAATGAG

TGAGCTAACTCACATTAATTGCGTTGCGCTCACTGCCCGCTTTCCAGTCGGGAAACCTGTCGTGCCAGCTGCATTAATGAATCGGCCAACGCGCGGGGAGAGGCGGTTTGCG
TATTGGGCGCTCTTCCGCTTCCTCGCTCACTGACTCGCTGCGCTCGGTCGTTCGGCTGCGGCGAGCGGTATCAGCTCACTCAAAGGCGGTAATACGGTTATCCACAGAATCA
GGGGATAACGCAGGAAAGAACATGTGAGCAAAAGGCCAGCAAAAGGCCAGGAACCGTAAAAAGGCCGCGTTGCTGGCGTTTTTCCATAGGCTCCGCCCCCCTGACGAGCATC
ACAAAAATCGACGCTCAAGTCAGAGGTGGCGAAACCCGACAGGACTATAAAGATACCAGGCGTTTCCCCCTGGAAGCTCCCTCGTGCGCTCTCCTGTTCCGACCCTGCCGCT
TACCGGATACCTGTCCGCCTTTCTCCCTTCGGGAAGCGTGGCGCTTTCTCAATGCTCACGCTGTAGGTATCTCAGTTCGGTGTAGGTCGTTCGCTCCAAGCTGGGCTGTGTG
CACGAACCCCCCGTTCAGCCCGACCGCTGCGCCTTATCCGGTAACTATCGTCTTGAGTCCAACCCGGTAAGACACGACTTATCGCCACTGGCAGCAGCCACTGGTAACAGGA
TTAGCAGAGCGAGGTATGTAGGCGGTGCTACAGAGTTCTTGAAGTGGTGGCCTAACTACGGCTACACTAGAAGGACAGTATTTGGTATCTGCGCTCTGCTGAAGCCAGTTAC
CTTCGGAAAAAGAGTTGGTAGCTCTTGATCCGGCAAACAAACCACCGCTGGTAGCGGTGGTTTTTTTGTTTGCAAGCAGCAGATTACGCGCAGAAAAAAAGGATCTCAAGAA
GATCCTTTGATCTTTTCTACGGGGTCTGACGCTCAGTGGAACGAAAACTCACGTTAAGGGATTTTGGTCATGAGATTATCAAAAAGGATCTTCACCTAGATCCTTTTAAATT
AAAAATGAAGTTTTAAATCAATCTAAAGTATATATGAGTAAACTTGGTCTGACAGTTACCAATGCTTAATCAGTGAGGCACCTATCTCAGCGATCTGTCTATTTCGTTCATC
CATAGTTGCCTGACTCCCCGTCGTGTAGATAACTACGATACGGGAGGGCTTACCATCTGGCCCCAGTGCTGCAATGATACCGCGAGACCCACGCTCACCGGCTCCAGATTTA
TCAGCAATAAACCAGCCAGCCGGAAGGGCCGAGCGCAGAAGTGGTCCTGCAACTTTATCCGCCTCCATCCAGTCTATTAATTGTTGCCGGGAAGCTAGAGTAAGTAGTTCGC
CAGTTAATAGTTTGCGCAACGTTGTTGCCATTGCTACAGGCATCGTGGTGTCACGCTCGTCGTTTGGTATGGCTTCATTCAGCTCCGGTTCCCAACGATCAAGGCGAGTTAC
ATGATCCCCCATGTTGTGCAAAAAAGCGGTTAGCTCCTTCGGTCCTCCGATCGTTGTCAGAAGTAAGTTGGCCGCAGTGTTATCACTCATGGTTATGGCAGCACTGCATAAT
TCTCTTACTGTCATGCCATCCGTAAGATGCTTTTCTGTGACTGGTGAGTACTCAACCAAGTCATTCTGAGAATAGTGTATGCGGCGACCGAGTTGCTCTTGCCCGGCGTCAA
TACGGGATAATACCGCGCCACATAGCAGAACTTTAAAAGTGCTCATCATTGGAAAACGTTCTTCGGGGCGAAAACTCTCAAGGATCTTACCGCTGTTGAGATCCAGTTCGAT
GTAACCCACTCGTGCACCCAACTGATCTTCAGCATCTTTTACTTTCACCAGCGTTTCTGGGTGAGCAAAAACAGGAAGGCAAAATGCCGCAAAAAAGGGAATAAGGGCGACA
CGGAAATGTTGAATACTCATACTCTTCCTTTTTCAATATTATTGAAGCATTTATCAGGGTTATTGTCTCATGAGCGGATACATATTTGAATGTATTTAGAAAAATAAACAAA
TAGGGGTTCCGCGCACATTTCCCCGAAAAGTGCCACCTGACGTC

#### EPG (I.p. 3F)-IRES-GCaMP6m:

GACGGATCGGGAGATCTCCCGATCCCCTATGGTCGACTCTCAGTACAATCTGCTCTGATGCCGCATAGTTAAGCCAGTATCTGCTCCCTGCTTGTGTGTTGGAGGTCGCTGA
GTAGTGCGCGAGCAAAATTTAAGCTACAACAAGGCAAGGCTTGACCGACAATTGCATGAAGAATCTGCTTAGGGTTAGGCGTTTTGCGCTGCTTCGCGATGTACGGGCCAGA
TATACGCGTTGACATTGATTATTGACTAGTTATTAATAGTAATCAATTACGGGGTCATTAGTTCATAGCCCATATATGGAGTTCCGCGTTACATAACTTACGGTAAATGGCC
CGCCTGGCTGACCGCCCAACGACCCCCGCCCATTGACGTCAATAATGACGTATGTTCCCATAGTAACGCCAATAGGGACTTTCCATTGACGTCAATGGGTGGACTATTTACG
GTAAACTGCCCACTTGGCAGTACATCAAGTGTATCATATGCCAAGTACGCCCCCTATTGACGTCAATGACGGTAAATGGCCCGCCTGGCATTATGCCCAGTACATGACCTTA
TGGGACTTTCCTACTTGGCAGTACATCTACGTATTAGTCATCGCTATTACCATGGTGATGCGGTTTTGGCAGTACATCAATGGGCGTGGATAGCGGTTTGACTCACGGGGAT
TTCCAAGTCTCCACCCCATTGACGTCAATGGGAGTTTGTTTTGGCACCAAAATCAACGGGACTTTCCAAAATGTCGTAACAACTCCGCCCCATTGACGCAAATGGGCGGTAG
GCGTGTACGGTGGGAGGTCTATATAAGCAGAGCTCTCTGGCTAACTAGAGAACCCACTGCTTACTGGCTTATCGAAATTAATACGACTCACTATAGGGAGACCCAAGCTGGC
TAGTTAAGCTTGGTACCGAGCTCGGATCCACTAGTCCAGTGTGGTGGAATTGCCCTTCCAGTACCCTTCACCATGAAGTGTGTACTTTTGGGATTCGCAGCAGTGATCGGAT
TCTTCGCGATCGCGGAGTCTCTTACCTGTAACACATGCTCAGTGAGTCTGATTGGAATATGTCTGAATCCCGCAACAGCGACTTGCTCCACCAACACATCCGTCTGCACCAC
AGGAAGAGCCAGTTTCACCAGCGGCAGCGGCCTGCTGAACTTCAACTCCCAGGGCTGCACGGAGGGAGCTCAGTGTAATGGCACCGTGTCCGGGTCCATCCTGGGTGCGTCG
TACACGGTCACTCAAACCTGCTGCAGCACAAACAACTGCAACCCCGTGACCAGCGGCGCCTCCTACGTCCAGATCTCCGTCAGCGCGGCCCTGAGCGCCGCCCTGCTGGCCT
GCGTCTGGGGCCAGTCCGTCTACGACTACAAGGATGACGATGACAAGGATTACAAAGACGACGATGATAAGGACTATAAGGATGATGACGACAAATAATAGCAATTCCTCGA
CGACTGCATAGGGTTACCCCCCTCTCCCTCCCCCCCCCCTAACGTTACTGGCCGAAGCCGCTTGGAATAAGGCCGGTGTGCGTTTGTCTATATGTTATTTTCCACCATATTG
CCGTCTTTTGGCAATGTGAGGGCCCGGAAACCTGGCCCTGTCTTCTTGACGAGCATTCCTAGGGGTCTTTCCCCTCTCGCCAAAGGAATGCAAGGTCTGTTGAATGTCGTGA
AGGAAGCAGTTCCTCTGGAAGCTTCTTGAAGACAAACAACGTCTGTAGCGACCCTTTGCAGGCAGCGGAACCCCCCACCTGGCGACAGGTGCCTCTGCGGCCAAAAGCCACG
TGTATAAGATACACCTGCAAAGGCGGCACAACCCCAGTGCCACGTTGTGAGTTGGATAGTTGTGGAAAGAGTCAAATGGCTCTCCTCAAGCGTATTCAACAAGGGGCTGAAG
GATGCCCAGAAGGTACCCCATTGTATGGGATCTGATCTGGGGCCTCGGTGCACATGCTTTACATGTGTTTAGTCGAGGTTAAAAAACGTCTAGGCCCCCCGAACCACGGGGA
CGTGGTTTTCCTTTGAAAAACACGATGATAATGGCCACAACCATGGGTTCTCATCATCATCATCATCATGGTATGGCTAGCATGACTGGTGGACAGCAAATGGGTCGGGATC
TGTACGACGATGACGATAAGGATCTCGCCACCATGGTCGACTCATCACGTCGTAAGTGGAATAAGACAGGTCACGCAGTCAGAGCTATAGGTCGGCTGAGCTCACTCGAGAA
CGTCTATATCAAGGCCGACAAGCAGAAGAACGGCATCAAGGCGAACTTCAAGATCCGCCACAACATCGAGGACGGCGGCGTGCAGCTCGCCTACCACTACCAGCAGAACACC
CCCATCGGCGACGGCCCCGTGCTGCTGCCCGACAACCACTACCTGAGCGTGCAGTCCAAACTTTCGAAAGACCCCAACGAGAAGCGCGATCACATGGTCCTGCTGGAGTTCG
TGACCGCCGCCGGGATCACTCTCGGCATGGACGAGCTGTACAAGGGCGGTACCGGAGGGAGCATGGTGAGCAAGGGCGAGGAGCTGTTCACCGGGGTGGTGCCCATCCTGGT
CGAGCTGGACGGCGACGTAAACGGCCACAAGTTCAGCGTGTCCGGCGAGGGTGAGGGCGATGCCACCTACGGCAAGCTGACCCTGAAGTTCATCTGCACCACCGGCAAGCTG
CCCGTGCCCTGGCCCACCCTCGTGACCACCCTGACCTACGGCGTGCAGTGCTTCAGCCGCTACCCCGACCACATGAAGCAGCACGACTTCTTCAAGTCCGCCATGCCCGAAG
GCTACATCCAGGAGCGCACCATCTTCTTCAAGGACGACGGCAACTACAAGACCCGCGCCGAGGTGAAGTTCGAGGGCGACACCCTGGTGAACCGCATCGAGCTGAAGGGCAT
CGACTTCAAGGAGGACGGCAACATCCTGGGGCACAAGCTGGAGTACAACCTGCCGGACCAACTGACTGAAGAGCAGATCGCAGAATTTAAAGAGGCTTTCTCCCTATTTGAC
AAGGACGGGGATGGGACAATAACAACCAAGGAGCTGGGGACGGTGATGCGGTCTCTGGGGCAGAACCCCACAGAAGCAGAGCTGCAGGACATGATCAATGAAGTAGATGCCG
ACGGTGACGGCACAATCGACTTCCCTGAGTTCCTGACAATGATGGCAAGAAAAGGGAGCTACAGGGACACGGAAGAAGAAATTAGAGAAGCGTTCGGTGTGTTTGATAAGGA
TGGCAATGGCTACATCAGTGCAGCAGAGCTTCGCCACGTGATGACAAACCTTGGAGAGAAGTTAACAGATGAAGAGGTTGATGAAATGATCAGGGAAGCAGACATCGATGGG
GATGGTCAGGTAAACTACGAAGAGTTTGTACAAATGATGACAGCGAAGTGAGAATTCTGCAGATATAAGGGCAATTCTGCAGATATCCAGCACAGTGGCGGCCGCTCGAGTC
TAGAGGGCCCGCGGTTCGAAGGTAAGCCTATCCCTAACCCTCTCCTCGGTCTCGATTCTACGCGTACCGGTCATCATCACCATCACCATTGAGTTTAAACCCGCTGATCAGC
CTCGACTGTGCCTTCTAGTTGCCAGCCATCTGTTGTTTGCCCCTCCCCCGTGCCTTCCTTGACCCTGGAAGGTGCCACTCCCACTGTCCTTTCCTAATAAAATGAGGAAATT
GCATCGCATTGTCTGAGTAGGTGTCATTCTATTCTGGGGGGTGGGGTGGGGCAGGACAGCAAGGGGGAGGATTGGGAAGACAATAGCAGGCATGCTGGGGATGCGGTGGGCT
CTATGGCTTCTGAGGCGGAAAGAACCAGCTGGGGCTCTAGGGGGTATCCCCACGCGCCCTGTAGCGGCGCATTAAGCGCGGCGGGTGTGGTGGTTACGCGCAGCGTGACCGC
TACACTTGCCAGCGCCCTAGCGCCCGCTCCTTTCGCTTTCTTCCCTTCCTTTCTCGCCACGTTCGCCGGCTTTCCCCGTCAAGCTCTAAATCGGGGCATCCCTTTAGGGTTC
CGATTTAGTGCTTTACGGCACCTCGACCCCAAAAAACTTGATTAGGGTGATGGTTCACGTAGTGGGCCATCGCCCTGATAGACGGTTTTTCGCCCTTTGACGTTGGAGTCCA
CGTTCTTTAATAGTGGACTCTTGTTCCAAACTGGAACAACACTCAACCCTATCTCGGTCTATTCTTTTGATTTATAAGGGATTTTGGGGATTTCGGCCTATTGGTTAAAAAA
TGAGCTGATTTAACAAAAATTTAACGCGAATTAATTCTGTGGAATGTGTGTCAGTTAGGGTGTGGAAAGTCCCCAGGCTCCCCAGGCAGGCAGAAGTATGCAAAGCATGCAT
CTCAATTAGTCAGCAACCAGGTGTGGAAAGTCCCCAGGCTCCCCAGCAGGCAGAAGTATGCAAAGCATGCATCTCAATTAGTCAGCAACCATAGTCCCGCCCCTAACTCCGC
CCATCCCGCCCCTAACTCCGCCCAGTTCCGCCCATTCTCCGCCCCATGGCTGACTAATTTTTTTTATTTATGCAGAGGCCGAGGCCGCCTCTGCCTCTGAGCTATTCCAGAA
GTAGTGAGGAGGCTTTTTTGGAGGCCTAGGCTTTTGCAAAAAGCTCCCGGGAGCTTGTATATCCATTTTCGGATCTGATCAAGAGACAGGATGAGGATCGTTTCGCATGATT
GAACAAGATGGATTGCACGCAGGTTCTCCGGCCGCTTGGGTGGAGAGGCTATTCGGCTATGACTGGGCACAACAGACAATCGGCTGCTCTGATGCCGCCGTGTTCCGGCTGT
CAGCGCAGGGGCGCCCGGTTCTTTTTGTCAAGACCGACCTGTCCGGTGCCCTGAATGAACTGCAGGACGAGGCAGCGCGGCTATCGTGGCTGGCCACGACGGGCGTTCCTTG
CGCAGCTGTGCTCGACGTTGTCACTGAAGCGGGAAGGGACTGGCTGCTATTGGGCGAAGTGCCGGGGCAGGATCTCCTGTCATCTCACCTTGCTCCTGCCGAGAAAGTATCC
ATCATGGCTGATGCAATGCGGCGGCTGCATACGCTTGATCCGGCTACCTGCCCATTCGACCACCAAGCGAAACATCGCATCGAGCGAGCACGTACTCGGATGGAAGCCGGTC
TTGTCGATCAGGATGATCTGGACGAAGAGCATCAGGGGCTCGCGCCAGCCGAACTGTTCGCCAGGCTCAAGGCGCGCATGCCCGACGGCGAGGATCTCGTCGTGACCCATGG
CGATGCCTGCTTGCCGAATATCATGGTGGAAAATGGCCGCTTTTCTGGATTCATCGACTGTGGCCGGCTGGGTGTGGCGGACCGCTATCAGGACATAGCGTTGGCTACCCGT
GATATTGCTGAAGAGCTTGGCGGCGAATGGGCTGACCGCTTCCTCGTGCTTTACGGTATCGCCGCTCCCGATTCGCAGCGCATCGCCTTCTATCGCCTTCTTGACGAGTTCT
TCTGAGCGGGACTCTGGGGTTCGCGAAATGACCGACCAAGCGACGCCCAACCTGCCATCACGAGATTTCGATTCCACCGCCGCCTTCTATGAAAGGTTGGGCTTCGGAATCG
TTTTCCGGGACGCCGGCTGGATGATCCTCCAGCGCGGGGATCTCATGCTGGAGTTCTTCGCCCACCCCAACTTGTTTATTGCAGCTTATAATGGTTACAAATAAAGCAATAG
CATCACAAATTTCACAAATAAAGCATTTTTTTCACTGCATTCTAGTTGTGGTTTGTCCAAACTCATCAATGTATCTTATCATGTCTGTATACCGTCGACCTCTAGCTAGAGC
TTGGCGTAATCATGGTCATAGCTGTTTCCTGTGTGAAATTGTTATCCGCTCACAATTCCACACAACATACGAGCCGGAAGCATAAAGTGTAAAGCCTGGGGTGCCTAATGAG
TGAGCTAACTCACATTAATTGCGTTGCGCTCACTGCCCGCTTTCCAGTCGGGAAACCTGTCGTGCCAGCTGCATTAATGAATCGGCCAACGCGCGGGGAGAGGCGGTTTGCG
TATTGGGCGCTCTTCCGCTTCCTCGCTCACTGACTCGCTGCGCTCGGTCGTTCGGCTGCGGCGAGCGGTATCAGCTCACTCAAAGGCGGTAATACGGTTATCCACAGAATCA
GGGGATAACGCAGGAAAGAACATGTGAGCAAAAGGCCAGCAAAAGGCCAGGAACCGTAAAAAGGCCGCGTTGCTGGCGTTTTTCCATAGGCTCCGCCCCCCTGACGAGCATC
ACAAAAATCGACGCTCAAGTCAGAGGTGGCGAAACCCGACAGGACTATAAAGATACCAGGCGTTTCCCCCTGGAAGCTCCCTCGTGCGCTCTCCTGTTCCGACCCTGCCGCT
TACCGGATACCTGTCCGCCTTTCTCCCTTCGGGAAGCGTGGCGCTTTCTCAATGCTCACGCTGTAGGTATCTCAGTTCGGTGTAGGTCGTTCGCTCCAAGCTGGGCTGTGTG

CACGAACCCCCCGTTCAGCCCGACCGCTGCGCCTTATCCGGTAACTATCGTCTTGAGTCCAACCCGGTAAGACACGACTTATCGCCACTGGCAGCAGCCACTGGTAACAGGA
TTAGCAGAGCGAGGTATGTAGGCGGTGCTACAGAGTTCTTGAAGTGGTGGCCTAACTACGGCTACACTAGAAGGACAGTATTTGGTATCTGCGCTCTGCTGAAGCCAGTTAC
CTTCGGAAAAAGAGTTGGTAGCTCTTGATCCGGCAAACAAACCACCGCTGGTAGCGGTGGTTTTTTTGTTTGCAAGCAGCAGATTACGCGCAGAAAAAAAGGATCTCAAGAA
GATCCTTTGATCTTTTCTACGGGGTCTGACGCTCAGTGGAACGAAAACTCACGTTAAGGGATTTTGGTCATGAGATTATCAAAAAGGATCTTCACCTAGATCCTTTTAAATT
AAAAATGAAGTTTTAAATCAATCTAAAGTATATATGAGTAAACTTGGTCTGACAGTTACCAATGCTTAATCAGTGAGGCACCTATCTCAGCGATCTGTCTATTTCGTTCATC
CATAGTTGCCTGACTCCCCGTCGTGTAGATAACTACGATACGGGAGGGCTTACCATCTGGCCCCAGTGCTGCAATGATACCGCGAGACCCACGCTCACCGGCTCCAGATTTA
TCAGCAATAAACCAGCCAGCCGGAAGGGCCGAGCGCAGAAGTGGTCCTGCAACTTTATCCGCCTCCATCCAGTCTATTAATTGTTGCCGGGAAGCTAGAGTAAGTAGTTCGC
CAGTTAATAGTTTGCGCAACGTTGTTGCCATTGCTACAGGCATCGTGGTGTCACGCTCGTCGTTTGGTATGGCTTCATTCAGCTCCGGTTCCCAACGATCAAGGCGAGTTAC
ATGATCCCCCATGTTGTGCAAAAAAGCGGTTAGCTCCTTCGGTCCTCCGATCGTTGTCAGAAGTAAGTTGGCCGCAGTGTTATCACTCATGGTTATGGCAGCACTGCATAAT
TCTCTTACTGTCATGCCATCCGTAAGATGCTTTTCTGTGACTGGTGAGTACTCAACCAAGTCATTCTGAGAATAGTGTATGCGGCGACCGAGTTGCTCTTGCCCGGCGTCAA
TACGGGATAATACCGCGCCACATAGCAGAACTTTAAAAGTGCTCATCATTGGAAAACGTTCTTCGGGGCGAAAACTCTCAAGGATCTTACCGCTGTTGAGATCCAGTTCGAT
GTAACCCACTCGTGCACCCAACTGATCTTCAGCATCTTTTACTTTCACCAGCGTTTCTGGGTGAGCAAAAACAGGAAGGCAAAATGCCGCAAAAAAGGGAATAAGGGCGACA
CGGAAATGTTGAATACTCATACTCTTCCTTTTTCAATATTATTGAAGCATTTATCAGGGTTATTGTCTCATGAGCGGATACATATTTGAATGTATTTAGAAAAATAAACAAA
TAGGGGTTCCGCGCACATTTCCCCGAAAAGTGCCACCTGACGTC

#### EPG (M.e. 3F)-IRES-GCaMP6m:

GACGGATCGGGAGATCTCCCGATCCCCTATGGTCGACTCTCAGTACAATCTGCTCTGATGCCGCATAGTTAAGCCAGTATCTGCTCCCTGCTTGTGTGTTGGAGGTCGCTGA
GTAGTGCGCGAGCAAAATTTAAGCTACAACAAGGCAAGGCTTGACCGACAATTGCATGAAGAATCTGCTTAGGGTTAGGCGTTTTGCGCTGCTTCGCGATGTACGGGCCAGA
TATACGCGTTGACATTGATTATTGACTAGTTATTAATAGTAATCAATTACGGGGTCATTAGTTCATAGCCCATATATGGAGTTCCGCGTTACATAACTTACGGTAAATGGCC
CGCCTGGCTGACCGCCCAACGACCCCCGCCCATTGACGTCAATAATGACGTATGTTCCCATAGTAACGCCAATAGGGACTTTCCATTGACGTCAATGGGTGGACTATTTACG
GTAAACTGCCCACTTGGCAGTACATCAAGTGTATCATATGCCAAGTACGCCCCCTATTGACGTCAATGACGGTAAATGGCCCGCCTGGCATTATGCCCAGTACATGACCTTA
TGGGACTTTCCTACTTGGCAGTACATCTACGTATTAGTCATCGCTATTACCATGGTGATGCGGTTTTGGCAGTACATCAATGGGCGTGGATAGCGGTTTGACTCACGGGGAT
TTCCAAGTCTCCACCCCATTGACGTCAATGGGAGTTTGTTTTGGCACCAAAATCAACGGGACTTTCCAAAATGTCGTAACAACTCCGCCCCATTGACGCAAATGGGCGGTAG
GCGTGTACGGTGGGAGGTCTATATAAGCAGAGCTCTCTGGCTAACTAGAGAACCCACTGCTTACTGGCTTATCGAAATTAATACGACTCACTATAGGGAGACCCAAGCTGGC
TAGTTAAGCTTGGTACCGAGCTCGGATCCACTAGTCCAGTGTGGTGGAATTGCCCTTCCAGTACCCTTCACCATGAAGTGTGTACTTTTGGGATTCGCAGCAGTGATCGGAT
TCTTCGCGATCGCGGAGTCTCTTACCTGTAACACATGCTCAGTGAGTCTGATTGGAATATGTCTGAATCCCGCAACAGCGACTTGCTCCACCAACACATCCGTCTGCACCAC
AGGAAGAGCCAGTTTCACCGGAATCACCGGCTTCCTGGGCTTCAACTCCCAGGGCTGCACGGAGGGAGCTCAGTGTAATGGCACCGTGTCCGGGTCCATCCTGGGTGCGTCG
TACACGGTCACTCAAACCTGCTGCAGCACAAACAACTGCAACCCCGTGACCAGCGGCGCCTCCTACGTCCAGATCTCCGTCAGCGCGGCCCTGAGCGCCGCCCTGCTGGCCT
GCGTCTGGGGCCAGTCCGTCTACGACTACAAGGATGACGATGACAAGGATTACAAAGACGACGATGATAAGGACTATAAGGATGATGACGACAAATAATAGCAATTCCTCGA
CGACTGCATAGGGTTACCCCCCTCTCCCTCCCCCCCCCCTAACGTTACTGGCCGAAGCCGCTTGGAATAAGGCCGGTGTGCGTTTGTCTATATGTTATTTTCCACCATATTG
CCGTCTTTTGGCAATGTGAGGGCCCGGAAACCTGGCCCTGTCTTCTTGACGAGCATTCCTAGGGGTCTTTCCCCTCTCGCCAAAGGAATGCAAGGTCTGTTGAATGTCGTGA
AGGAAGCAGTTCCTCTGGAAGCTTCTTGAAGACAAACAACGTCTGTAGCGACCCTTTGCAGGCAGCGGAACCCCCCACCTGGCGACAGGTGCCTCTGCGGCCAAAAGCCACG
TGTATAAGATACACCTGCAAAGGCGGCACAACCCCAGTGCCACGTTGTGAGTTGGATAGTTGTGGAAAGAGTCAAATGGCTCTCCTCAAGCGTATTCAACAAGGGGCTGAAG
GATGCCCAGAAGGTACCCCATTGTATGGGATCTGATCTGGGGCCTCGGTGCACATGCTTTACATGTGTTTAGTCGAGGTTAAAAAACGTCTAGGCCCCCCGAACCACGGGGA
CGTGGTTTTCCTTTGAAAAACACGATGATAATGGCCACAACCATGGGTTCTCATCATCATCATCATCATGGTATGGCTAGCATGACTGGTGGACAGCAAATGGGTCGGGATC
TGTACGACGATGACGATAAGGATCTCGCCACCATGGTCGACTCATCACGTCGTAAGTGGAATAAGACAGGTCACGCAGTCAGAGCTATAGGTCGGCTGAGCTCACTCGAGAA
CGTCTATATCAAGGCCGACAAGCAGAAGAACGGCATCAAGGCGAACTTCAAGATCCGCCACAACATCGAGGACGGCGGCGTGCAGCTCGCCTACCACTACCAGCAGAACACC
CCCATCGGCGACGGCCCCGTGCTGCTGCCCGACAACCACTACCTGAGCGTGCAGTCCAAACTTTCGAAAGACCCCAACGAGAAGCGCGATCACATGGTCCTGCTGGAGTTCG
TGACCGCCGCCGGGATCACTCTCGGCATGGACGAGCTGTACAAGGGCGGTACCGGAGGGAGCATGGTGAGCAAGGGCGAGGAGCTGTTCACCGGGGTGGTGCCCATCCTGGT
CGAGCTGGACGGCGACGTAAACGGCCACAAGTTCAGCGTGTCCGGCGAGGGTGAGGGCGATGCCACCTACGGCAAGCTGACCCTGAAGTTCATCTGCACCACCGGCAAGCTG
CCCGTGCCCTGGCCCACCCTCGTGACCACCCTGACCTACGGCGTGCAGTGCTTCAGCCGCTACCCCGACCACATGAAGCAGCACGACTTCTTCAAGTCCGCCATGCCCGAAG
GCTACATCCAGGAGCGCACCATCTTCTTCAAGGACGACGGCAACTACAAGACCCGCGCCGAGGTGAAGTTCGAGGGCGACACCCTGGTGAACCGCATCGAGCTGAAGGGCAT
CGACTTCAAGGAGGACGGCAACATCCTGGGGCACAAGCTGGAGTACAACCTGCCGGACCAACTGACTGAAGAGCAGATCGCAGAATTTAAAGAGGCTTTCTCCCTATTTGAC
AAGGACGGGGATGGGACAATAACAACCAAGGAGCTGGGGACGGTGATGCGGTCTCTGGGGCAGAACCCCACAGAAGCAGAGCTGCAGGACATGATCAATGAAGTAGATGCCG
ACGGTGACGGCACAATCGACTTCCCTGAGTTCCTGACAATGATGGCAAGAAAAGGGAGCTACAGGGACACGGAAGAAGAAATTAGAGAAGCGTTCGGTGTGTTTGATAAGGA
TGGCAATGGCTACATCAGTGCAGCAGAGCTTCGCCACGTGATGACAAACCTTGGAGAGAAGTTAACAGATGAAGAGGTTGATGAAATGATCAGGGAAGCAGACATCGATGGG
GATGGTCAGGTAAACTACGAAGAGTTTGTACAAATGATGACAGCGAAGTGAGAATTCTGCAGATATAAGGGCAATTCTGCAGATATCCAGCACAGTGGCGGCCGCTCGAGTC
TAGAGGGCCCGCGGTTCGAAGGTAAGCCTATCCCTAACCCTCTCCTCGGTCTCGATTCTACGCGTACCGGTCATCATCACCATCACCATTGAGTTTAAACCCGCTGATCAGC
CTCGACTGTGCCTTCTAGTTGCCAGCCATCTGTTGTTTGCCCCTCCCCCGTGCCTTCCTTGACCCTGGAAGGTGCCACTCCCACTGTCCTTTCCTAATAAAATGAGGAAATT
GCATCGCATTGTCTGAGTAGGTGTCATTCTATTCTGGGGGGTGGGGTGGGGCAGGACAGCAAGGGGGAGGATTGGGAAGACAATAGCAGGCATGCTGGGGATGCGGTGGGCT
CTATGGCTTCTGAGGCGGAAAGAACCAGCTGGGGCTCTAGGGGGTATCCCCACGCGCCCTGTAGCGGCGCATTAAGCGCGGCGGGTGTGGTGGTTACGCGCAGCGTGACCGC
TACACTTGCCAGCGCCCTAGCGCCCGCTCCTTTCGCTTTCTTCCCTTCCTTTCTCGCCACGTTCGCCGGCTTTCCCCGTCAAGCTCTAAATCGGGGCATCCCTTTAGGGTTC
CGATTTAGTGCTTTACGGCACCTCGACCCCAAAAAACTTGATTAGGGTGATGGTTCACGTAGTGGGCCATCGCCCTGATAGACGGTTTTTCGCCCTTTGACGTTGGAGTCCA
CGTTCTTTAATAGTGGACTCTTGTTCCAAACTGGAACAACACTCAACCCTATCTCGGTCTATTCTTTTGATTTATAAGGGATTTTGGGGATTTCGGCCTATTGGTTAAAAAA
TGAGCTGATTTAACAAAAATTTAACGCGAATTAATTCTGTGGAATGTGTGTCAGTTAGGGTGTGGAAAGTCCCCAGGCTCCCCAGGCAGGCAGAAGTATGCAAAGCATGCAT
CTCAATTAGTCAGCAACCAGGTGTGGAAAGTCCCCAGGCTCCCCAGCAGGCAGAAGTATGCAAAGCATGCATCTCAATTAGTCAGCAACCATAGTCCCGCCCCTAACTCCGC
CCATCCCGCCCCTAACTCCGCCCAGTTCCGCCCATTCTCCGCCCCATGGCTGACTAATTTTTTTTATTTATGCAGAGGCCGAGGCCGCCTCTGCCTCTGAGCTATTCCAGAA
GTAGTGAGGAGGCTTTTTTGGAGGCCTAGGCTTTTGCAAAAAGCTCCCGGGAGCTTGTATATCCATTTTCGGATCTGATCAAGAGACAGGATGAGGATCGTTTCGCATGATT
GAACAAGATGGATTGCACGCAGGTTCTCCGGCCGCTTGGGTGGAGAGGCTATTCGGCTATGACTGGGCACAACAGACAATCGGCTGCTCTGATGCCGCCGTGTTCCGGCTGT
CAGCGCAGGGGCGCCCGGTTCTTTTTGTCAAGACCGACCTGTCCGGTGCCCTGAATGAACTGCAGGACGAGGCAGCGCGGCTATCGTGGCTGGCCACGACGGGCGTTCCTTG
CGCAGCTGTGCTCGACGTTGTCACTGAAGCGGGAAGGGACTGGCTGCTATTGGGCGAAGTGCCGGGGCAGGATCTCCTGTCATCTCACCTTGCTCCTGCCGAGAAAGTATCC
ATCATGGCTGATGCAATGCGGCGGCTGCATACGCTTGATCCGGCTACCTGCCCATTCGACCACCAAGCGAAACATCGCATCGAGCGAGCACGTACTCGGATGGAAGCCGGTC
TTGTCGATCAGGATGATCTGGACGAAGAGCATCAGGGGCTCGCGCCAGCCGAACTGTTCGCCAGGCTCAAGGCGCGCATGCCCGACGGCGAGGATCTCGTCGTGACCCATGG
CGATGCCTGCTTGCCGAATATCATGGTGGAAAATGGCCGCTTTTCTGGATTCATCGACTGTGGCCGGCTGGGTGTGGCGGACCGCTATCAGGACATAGCGTTGGCTACCCGT
GATATTGCTGAAGAGCTTGGCGGCGAATGGGCTGACCGCTTCCTCGTGCTTTACGGTATCGCCGCTCCCGATTCGCAGCGCATCGCCTTCTATCGCCTTCTTGACGAGTTCT
TCTGAGCGGGACTCTGGGGTTCGCGAAATGACCGACCAAGCGACGCCCAACCTGCCATCACGAGATTTCGATTCCACCGCCGCCTTCTATGAAAGGTTGGGCTTCGGAATCG
TTTTCCGGGACGCCGGCTGGATGATCCTCCAGCGCGGGGATCTCATGCTGGAGTTCTTCGCCCACCCCAACTTGTTTATTGCAGCTTATAATGGTTACAAATAAAGCAATAG
CATCACAAATTTCACAAATAAAGCATTTTTTTCACTGCATTCTAGTTGTGGTTTGTCCAAACTCATCAATGTATCTTATCATGTCTGTATACCGTCGACCTCTAGCTAGAGC
TTGGCGTAATCATGGTCATAGCTGTTTCCTGTGTGAAATTGTTATCCGCTCACAATTCCACACAACATACGAGCCGGAAGCATAAAGTGTAAAGCCTGGGGTGCCTAATGAG
TGAGCTAACTCACATTAATTGCGTTGCGCTCACTGCCCGCTTTCCAGTCGGGAAACCTGTCGTGCCAGCTGCATTAATGAATCGGCCAACGCGCGGGGAGAGGCGGTTTGCG
TATTGGGCGCTCTTCCGCTTCCTCGCTCACTGACTCGCTGCGCTCGGTCGTTCGGCTGCGGCGAGCGGTATCAGCTCACTCAAAGGCGGTAATACGGTTATCCACAGAATCA
GGGGATAACGCAGGAAAGAACATGTGAGCAAAAGGCCAGCAAAAGGCCAGGAACCGTAAAAAGGCCGCGTTGCTGGCGTTTTTCCATAGGCTCCGCCCCCCTGACGAGCATC
ACAAAAATCGACGCTCAAGTCAGAGGTGGCGAAACCCGACAGGACTATAAAGATACCAGGCGTTTCCCCCTGGAAGCTCCCTCGTGCGCTCTCCTGTTCCGACCCTGCCGCT
TACCGGATACCTGTCCGCCTTTCTCCCTTCGGGAAGCGTGGCGCTTTCTCAATGCTCACGCTGTAGGTATCTCAGTTCGGTGTAGGTCGTTCGCTCCAAGCTGGGCTGTGTG
CACGAACCCCCCGTTCAGCCCGACCGCTGCGCCTTATCCGGTAACTATCGTCTTGAGTCCAACCCGGTAAGACACGACTTATCGCCACTGGCAGCAGCCACTGGTAACAGGA
TTAGCAGAGCGAGGTATGTAGGCGGTGCTACAGAGTTCTTGAAGTGGTGGCCTAACTACGGCTACACTAGAAGGACAGTATTTGGTATCTGCGCTCTGCTGAAGCCAGTTAC
CTTCGGAAAAAGAGTTGGTAGCTCTTGATCCGGCAAACAAACCACCGCTGGTAGCGGTGGTTTTTTTGTTTGCAAGCAGCAGATTACGCGCAGAAAAAAAGGATCTCAAGAA
GATCCTTTGATCTTTTCTACGGGGTCTGACGCTCAGTGGAACGAAAACTCACGTTAAGGGATTTTGGTCATGAGATTATCAAAAAGGATCTTCACCTAGATCCTTTTAAATT
AAAAATGAAGTTTTAAATCAATCTAAAGTATATATGAGTAAACTTGGTCTGACAGTTACCAATGCTTAATCAGTGAGGCACCTATCTCAGCGATCTGTCTATTTCGTTCATC

CATAGTTGCCTGACTCCCCGTCGTGTAGATAACTACGATACGGGAGGGCTTACCATCTGGCCCCAGTGCTGCAATGATACCGCGAGACCCACGCTCACCGGCTCCAGATTTA
TCAGCAATAAACCAGCCAGCCGGAAGGGCCGAGCGCAGAAGTGGTCCTGCAACTTTATCCGCCTCCATCCAGTCTATTAATTGTTGCCGGGAAGCTAGAGTAAGTAGTTCGC
CAGTTAATAGTTTGCGCAACGTTGTTGCCATTGCTACAGGCATCGTGGTGTCACGCTCGTCGTTTGGTATGGCTTCATTCAGCTCCGGTTCCCAACGATCAAGGCGAGTTAC
ATGATCCCCCATGTTGTGCAAAAAAGCGGTTAGCTCCTTCGGTCCTCCGATCGTTGTCAGAAGTAAGTTGGCCGCAGTGTTATCACTCATGGTTATGGCAGCACTGCATAAT
TCTCTTACTGTCATGCCATCCGTAAGATGCTTTTCTGTGACTGGTGAGTACTCAACCAAGTCATTCTGAGAATAGTGTATGCGGCGACCGAGTTGCTCTTGCCCGGCGTCAA
TACGGGATAATACCGCGCCACATAGCAGAACTTTAAAAGTGCTCATCATTGGAAAACGTTCTTCGGGGCGAAAACTCTCAAGGATCTTACCGCTGTTGAGATCCAGTTCGAT
GTAACCCACTCGTGCACCCAACTGATCTTCAGCATCTTTTACTTTCACCAGCGTTTCTGGGTGAGCAAAAACAGGAAGGCAAAATGCCGCAAAAAAGGGAATAAGGGCGACA
CGGAAATGTTGAATACTCATACTCTTCCTTTTTCAATATTATTGAAGCATTTATCAGGGTTATTGTCTCATGAGCGGATACATATTTGAATGTATTTAGAAAAATAAACAAA
TAGGGGTTCCGCGCACATTTCCCCGAAAAGTGCCACCTGACGTC

#### EPG (S.m. 3F)-IRES-GCaMP6m:

GACGGATCGGGAGATCTCCCGATCCCCTATGGTCGACTCTCAGTACAATCTGCTCTGATGCCGCATAGTTAAGCCAGTATCTGCTCCCTGCTTGTGTGTTGGAGGTCGCTGA
GTAGTGCGCGAGCAAAATTTAAGCTACAACAAGGCAAGGCTTGACCGACAATTGCATGAAGAATCTGCTTAGGGTTAGGCGTTTTGCGCTGCTTCGCGATGTACGGGCCAGA
TATACGCGTTGACATTGATTATTGACTAGTTATTAATAGTAATCAATTACGGGGTCATTAGTTCATAGCCCATATATGGAGTTCCGCGTTACATAACTTACGGTAAATGGCC
CGCCTGGCTGACCGCCCAACGACCCCCGCCCATTGACGTCAATAATGACGTATGTTCCCATAGTAACGCCAATAGGGACTTTCCATTGACGTCAATGGGTGGACTATTTACG
GTAAACTGCCCACTTGGCAGTACATCAAGTGTATCATATGCCAAGTACGCCCCCTATTGACGTCAATGACGGTAAATGGCCCGCCTGGCATTATGCCCAGTACATGACCTTA
TGGGACTTTCCTACTTGGCAGTACATCTACGTATTAGTCATCGCTATTACCATGGTGATGCGGTTTTGGCAGTACATCAATGGGCGTGGATAGCGGTTTGACTCACGGGGAT
TTCCAAGTCTCCACCCCATTGACGTCAATGGGAGTTTGTTTTGGCACCAAAATCAACGGGACTTTCCAAAATGTCGTAACAACTCCGCCCCATTGACGCAAATGGGCGGTAG
GCGTGTACGGTGGGAGGTCTATATAAGCAGAGCTCTCTGGCTAACTAGAGAACCCACTGCTTACTGGCTTATCGAAATTAATACGACTCACTATAGGGAGACCCAAGCTGGC
TAGTTAAGCTTGGTACCGAGCTCGGATCCACTAGTCCAGTGTGGTGGAATTGCCCTTCCAGTACCCTTCACCATGAAGTGTGTACTTTTGGGATTCGCAGCAGTGATCGGAT
TCTTCGCGATCGCGGAGTCTCTTACCTGTAACACATGCTCAGTGAGTCTGATTGGAATATGTCTGAATCCCGCAACAGCGACTTGCTCCACCAACACATCCGTCTGCACCAC
AGGAAGAGCCAGTTTCCCCTCAATCACCGGCTTCCTGGGCTTCAACTCCCAGGGCTGCACGGAGGGAGCTCAGTGTAATGGCACCGTGTCCGGGTCCATCCTGGGTGCGTCG
TACACGGTCACTCAAACCTGCTGCAGCACAAACAACTGCAACCCCGTGACCAGCGGCGCCTCCTACGTCCAGATCTCCGTCAGCGCGGCCCTGAGCGCCGCCCTGCTGGCCT
GCGTCTGGGGCCAGTCCGTCTACGACTACAAGGATGACGATGACAAGGATTACAAAGACGACGATGATAAGGACTATAAGGATGATGACGACAAATAATAGCAATTCCTCGA
CGACTGCATAGGGTTACCCCCCTCTCCCTCCCCCCCCCCTAACGTTACTGGCCGAAGCCGCTTGGAATAAGGCCGGTGTGCGTTTGTCTATATGTTATTTTCCACCATATTG
CCGTCTTTTGGCAATGTGAGGGCCCGGAAACCTGGCCCTGTCTTCTTGACGAGCATTCCTAGGGGTCTTTCCCCTCTCGCCAAAGGAATGCAAGGTCTGTTGAATGTCGTGA
AGGAAGCAGTTCCTCTGGAAGCTTCTTGAAGACAAACAACGTCTGTAGCGACCCTTTGCAGGCAGCGGAACCCCCCACCTGGCGACAGGTGCCTCTGCGGCCAAAAGCCACG
TGTATAAGATACACCTGCAAAGGCGGCACAACCCCAGTGCCACGTTGTGAGTTGGATAGTTGTGGAAAGAGTCAAATGGCTCTCCTCAAGCGTATTCAACAAGGGGCTGAAG
GATGCCCAGAAGGTACCCCATTGTATGGGATCTGATCTGGGGCCTCGGTGCACATGCTTTACATGTGTTTAGTCGAGGTTAAAAAACGTCTAGGCCCCCCGAACCACGGGGA
CGTGGTTTTCCTTTGAAAAACACGATGATAATGGCCACAACCATGGGTTCTCATCATCATCATCATCATGGTATGGCTAGCATGACTGGTGGACAGCAAATGGGTCGGGATC
TGTACGACGATGACGATAAGGATCTCGCCACCATGGTCGACTCATCACGTCGTAAGTGGAATAAGACAGGTCACGCAGTCAGAGCTATAGGTCGGCTGAGCTCACTCGAGAA
CGTCTATATCAAGGCCGACAAGCAGAAGAACGGCATCAAGGCGAACTTCAAGATCCGCCACAACATCGAGGACGGCGGCGTGCAGCTCGCCTACCACTACCAGCAGAACACC
CCCATCGGCGACGGCCCCGTGCTGCTGCCCGACAACCACTACCTGAGCGTGCAGTCCAAACTTTCGAAAGACCCCAACGAGAAGCGCGATCACATGGTCCTGCTGGAGTTCG
TGACCGCCGCCGGGATCACTCTCGGCATGGACGAGCTGTACAAGGGCGGTACCGGAGGGAGCATGGTGAGCAAGGGCGAGGAGCTGTTCACCGGGGTGGTGCCCATCCTGGT
CGAGCTGGACGGCGACGTAAACGGCCACAAGTTCAGCGTGTCCGGCGAGGGTGAGGGCGATGCCACCTACGGCAAGCTGACCCTGAAGTTCATCTGCACCACCGGCAAGCTG
CCCGTGCCCTGGCCCACCCTCGTGACCACCCTGACCTACGGCGTGCAGTGCTTCAGCCGCTACCCCGACCACATGAAGCAGCACGACTTCTTCAAGTCCGCCATGCCCGAAG
GCTACATCCAGGAGCGCACCATCTTCTTCAAGGACGACGGCAACTACAAGACCCGCGCCGAGGTGAAGTTCGAGGGCGACACCCTGGTGAACCGCATCGAGCTGAAGGGCAT
CGACTTCAAGGAGGACGGCAACATCCTGGGGCACAAGCTGGAGTACAACCTGCCGGACCAACTGACTGAAGAGCAGATCGCAGAATTTAAAGAGGCTTTCTCCCTATTTGAC
AAGGACGGGGATGGGACAATAACAACCAAGGAGCTGGGGACGGTGATGCGGTCTCTGGGGCAGAACCCCACAGAAGCAGAGCTGCAGGACATGATCAATGAAGTAGATGCCG
ACGGTGACGGCACAATCGACTTCCCTGAGTTCCTGACAATGATGGCAAGAAAAGGGAGCTACAGGGACACGGAAGAAGAAATTAGAGAAGCGTTCGGTGTGTTTGATAAGGA
TGGCAATGGCTACATCAGTGCAGCAGAGCTTCGCCACGTGATGACAAACCTTGGAGAGAAGTTAACAGATGAAGAGGTTGATGAAATGATCAGGGAAGCAGACATCGATGGG
GATGGTCAGGTAAACTACGAAGAGTTTGTACAAATGATGACAGCGAAGTGAGAATTCTGCAGATATAAGGGCAATTCTGCAGATATCCAGCACAGTGGCGGCCGCTCGAGTC
TAGAGGGCCCGCGGTTCGAAGGTAAGCCTATCCCTAACCCTCTCCTCGGTCTCGATTCTACGCGTACCGGTCATCATCACCATCACCATTGAGTTTAAACCCGCTGATCAGC
CTCGACTGTGCCTTCTAGTTGCCAGCCATCTGTTGTTTGCCCCTCCCCCGTGCCTTCCTTGACCCTGGAAGGTGCCACTCCCACTGTCCTTTCCTAATAAAATGAGGAAATT
GCATCGCATTGTCTGAGTAGGTGTCATTCTATTCTGGGGGGTGGGGTGGGGCAGGACAGCAAGGGGGAGGATTGGGAAGACAATAGCAGGCATGCTGGGGATGCGGTGGGCT
CTATGGCTTCTGAGGCGGAAAGAACCAGCTGGGGCTCTAGGGGGTATCCCCACGCGCCCTGTAGCGGCGCATTAAGCGCGGCGGGTGTGGTGGTTACGCGCAGCGTGACCGC
TACACTTGCCAGCGCCCTAGCGCCCGCTCCTTTCGCTTTCTTCCCTTCCTTTCTCGCCACGTTCGCCGGCTTTCCCCGTCAAGCTCTAAATCGGGGCATCCCTTTAGGGTTC
CGATTTAGTGCTTTACGGCACCTCGACCCCAAAAAACTTGATTAGGGTGATGGTTCACGTAGTGGGCCATCGCCCTGATAGACGGTTTTTCGCCCTTTGACGTTGGAGTCCA
CGTTCTTTAATAGTGGACTCTTGTTCCAAACTGGAACAACACTCAACCCTATCTCGGTCTATTCTTTTGATTTATAAGGGATTTTGGGGATTTCGGCCTATTGGTTAAAAAA
TGAGCTGATTTAACAAAAATTTAACGCGAATTAATTCTGTGGAATGTGTGTCAGTTAGGGTGTGGAAAGTCCCCAGGCTCCCCAGGCAGGCAGAAGTATGCAAAGCATGCAT
CTCAATTAGTCAGCAACCAGGTGTGGAAAGTCCCCAGGCTCCCCAGCAGGCAGAAGTATGCAAAGCATGCATCTCAATTAGTCAGCAACCATAGTCCCGCCCCTAACTCCGC
CCATCCCGCCCCTAACTCCGCCCAGTTCCGCCCATTCTCCGCCCCATGGCTGACTAATTTTTTTTATTTATGCAGAGGCCGAGGCCGCCTCTGCCTCTGAGCTATTCCAGAA
GTAGTGAGGAGGCTTTTTTGGAGGCCTAGGCTTTTGCAAAAAGCTCCCGGGAGCTTGTATATCCATTTTCGGATCTGATCAAGAGACAGGATGAGGATCGTTTCGCATGATT
GAACAAGATGGATTGCACGCAGGTTCTCCGGCCGCTTGGGTGGAGAGGCTATTCGGCTATGACTGGGCACAACAGACAATCGGCTGCTCTGATGCCGCCGTGTTCCGGCTGT
CAGCGCAGGGGCGCCCGGTTCTTTTTGTCAAGACCGACCTGTCCGGTGCCCTGAATGAACTGCAGGACGAGGCAGCGCGGCTATCGTGGCTGGCCACGACGGGCGTTCCTTG
CGCAGCTGTGCTCGACGTTGTCACTGAAGCGGGAAGGGACTGGCTGCTATTGGGCGAAGTGCCGGGGCAGGATCTCCTGTCATCTCACCTTGCTCCTGCCGAGAAAGTATCC
ATCATGGCTGATGCAATGCGGCGGCTGCATACGCTTGATCCGGCTACCTGCCCATTCGACCACCAAGCGAAACATCGCATCGAGCGAGCACGTACTCGGATGGAAGCCGGTC
TTGTCGATCAGGATGATCTGGACGAAGAGCATCAGGGGCTCGCGCCAGCCGAACTGTTCGCCAGGCTCAAGGCGCGCATGCCCGACGGCGAGGATCTCGTCGTGACCCATGG
CGATGCCTGCTTGCCGAATATCATGGTGGAAAATGGCCGCTTTTCTGGATTCATCGACTGTGGCCGGCTGGGTGTGGCGGACCGCTATCAGGACATAGCGTTGGCTACCCGT
GATATTGCTGAAGAGCTTGGCGGCGAATGGGCTGACCGCTTCCTCGTGCTTTACGGTATCGCCGCTCCCGATTCGCAGCGCATCGCCTTCTATCGCCTTCTTGACGAGTTCT
TCTGAGCGGGACTCTGGGGTTCGCGAAATGACCGACCAAGCGACGCCCAACCTGCCATCACGAGATTTCGATTCCACCGCCGCCTTCTATGAAAGGTTGGGCTTCGGAATCG
TTTTCCGGGACGCCGGCTGGATGATCCTCCAGCGCGGGGATCTCATGCTGGAGTTCTTCGCCCACCCCAACTTGTTTATTGCAGCTTATAATGGTTACAAATAAAGCAATAG
CATCACAAATTTCACAAATAAAGCATTTTTTTCACTGCATTCTAGTTGTGGTTTGTCCAAACTCATCAATGTATCTTATCATGTCTGTATACCGTCGACCTCTAGCTAGAGC
TTGGCGTAATCATGGTCATAGCTGTTTCCTGTGTGAAATTGTTATCCGCTCACAATTCCACACAACATACGAGCCGGAAGCATAAAGTGTAAAGCCTGGGGTGCCTAATGAG
TGAGCTAACTCACATTAATTGCGTTGCGCTCACTGCCCGCTTTCCAGTCGGGAAACCTGTCGTGCCAGCTGCATTAATGAATCGGCCAACGCGCGGGGAGAGGCGGTTTGCG
TATTGGGCGCTCTTCCGCTTCCTCGCTCACTGACTCGCTGCGCTCGGTCGTTCGGCTGCGGCGAGCGGTATCAGCTCACTCAAAGGCGGTAATACGGTTATCCACAGAATCA
GGGGATAACGCAGGAAAGAACATGTGAGCAAAAGGCCAGCAAAAGGCCAGGAACCGTAAAAAGGCCGCGTTGCTGGCGTTTTTCCATAGGCTCCGCCCCCCTGACGAGCATC
ACAAAAATCGACGCTCAAGTCAGAGGTGGCGAAACCCGACAGGACTATAAAGATACCAGGCGTTTCCCCCTGGAAGCTCCCTCGTGCGCTCTCCTGTTCCGACCCTGCCGCT
TACCGGATACCTGTCCGCCTTTCTCCCTTCGGGAAGCGTGGCGCTTTCTCAATGCTCACGCTGTAGGTATCTCAGTTCGGTGTAGGTCGTTCGCTCCAAGCTGGGCTGTGTG
CACGAACCCCCCGTTCAGCCCGACCGCTGCGCCTTATCCGGTAACTATCGTCTTGAGTCCAACCCGGTAAGACACGACTTATCGCCACTGGCAGCAGCCACTGGTAACAGGA
TTAGCAGAGCGAGGTATGTAGGCGGTGCTACAGAGTTCTTGAAGTGGTGGCCTAACTACGGCTACACTAGAAGGACAGTATTTGGTATCTGCGCTCTGCTGAAGCCAGTTAC
CTTCGGAAAAAGAGTTGGTAGCTCTTGATCCGGCAAACAAACCACCGCTGGTAGCGGTGGTTTTTTTGTTTGCAAGCAGCAGATTACGCGCAGAAAAAAAGGATCTCAAGAA
GATCCTTTGATCTTTTCTACGGGGTCTGACGCTCAGTGGAACGAAAACTCACGTTAAGGGATTTTGGTCATGAGATTATCAAAAAGGATCTTCACCTAGATCCTTTTAAATT
AAAAATGAAGTTTTAAATCAATCTAAAGTATATATGAGTAAACTTGGTCTGACAGTTACCAATGCTTAATCAGTGAGGCACCTATCTCAGCGATCTGTCTATTTCGTTCATC
CATAGTTGCCTGACTCCCCGTCGTGTAGATAACTACGATACGGGAGGGCTTACCATCTGGCCCCAGTGCTGCAATGATACCGCGAGACCCACGCTCACCGGCTCCAGATTTA
TCAGCAATAAACCAGCCAGCCGGAAGGGCCGAGCGCAGAAGTGGTCCTGCAACTTTATCCGCCTCCATCCAGTCTATTAATTGTTGCCGGGAAGCTAGAGTAAGTAGTTCGC
CAGTTAATAGTTTGCGCAACGTTGTTGCCATTGCTACAGGCATCGTGGTGTCACGCTCGTCGTTTGGTATGGCTTCATTCAGCTCCGGTTCCCAACGATCAAGGCGAGTTAC
ATGATCCCCCATGTTGTGCAAAAAAGCGGTTAGCTCCTTCGGTCCTCCGATCGTTGTCAGAAGTAAGTTGGCCGCAGTGTTATCACTCATGGTTATGGCAGCACTGCATAAT
TCTCTTACTGTCATGCCATCCGTAAGATGCTTTTCTGTGACTGGTGAGTACTCAACCAAGTCATTCTGAGAATAGTGTATGCGGCGACCGAGTTGCTCTTGCCCGGCGTCAA

TACGGGATAATACCGCGCCACATAGCAGAACTTTAAAAGTGCTCATCATTGGAAAACGTTCTTCGGGGCGAAAACTCTCAAGGATCTTACCGCTGTTGAGATCCAGTTCGAT
GTAACCCACTCGTGCACCCAACTGATCTTCAGCATCTTTTACTTTCACCAGCGTTTCTGGGTGAGCAAAAACAGGAAGGCAAAATGCCGCAAAAAAGGGAATAAGGGCGACA
CGGAAATGTTGAATACTCATACTCTTCCTTTTTCAATATTATTGAAGCATTTATCAGGGTTATTGTCTCATGAGCGGATACATATTTGAATGTATTTAGAAAAATAAACAAA
TAGGGGTTCCGCGCACATTTCCCCGAAAAGTGCCACCTGACGTC
